# Proton-gated coincidence detection is a common feature of GPCR signaling

**DOI:** 10.1101/2021.01.08.425945

**Authors:** Nicholas J. Kapolka, Jacob B. Rowe, Geoffrey J. Taghon, William M. Morgan, Corin R. O’Shea, Daniel G. Isom

## Abstract

The evolutionary expansion of G protein-coupled receptors (GPCRs) has produced a rich diversity of transmembrane sensors for many physical and chemical signals. In humans alone, over 800 GPCRs detect stimuli such as light, hormones, and metabolites to guide cellular decision making primarily using intracellular G protein signaling networks. This diversity is further enriched by GPCRs that function as molecular logic gates capable of discerning multiple inputs to transduce cues encoded in complex, context-dependent signals. Here, we show that many GPCRs are switch-like Boolean-gated coincidence detectors that link proton (H^+^) binding to GPCR signaling. Using a panel of 28 receptors, covering 280 individual GPCR-Gα coupling combinations, we show that H^+^ gating both positively and negatively modulates and controls GPCR signaling. Notably, these observations extend to all modes of GPCR pharmacology including ligand efficacy, potency, and cooperativity. Additionally, we show that GPCR antagonism and constitutive activity are regulated by H^+^ gating and report the discovery of a new acid sensor, the adenosine A2a receptor (ADORA2A), which can be activated solely by acidic pH. Together, these findings establish a new paradigm for GPCR biology and pharmacology in acidified microenvironments such as endosomes, synapses, tumors, and ischemic vasculature.

## Introduction

Over hundreds of millions of years, eukaryotes have evolved vast sensor arrays of G protein-coupled receptors (GPCRs) (1). The size and functionality of this receptor superfamily stems primarily from the structural adaptability of its 7-transmembrane (7TM) core. The versatility of the 7TM architecture has facilitated an extensive diversification and expansion of species-specific GPCR repertoires that can detect thousands of physical and chemical signals. In humans, more than 800 GPCRs detect inputs such as light, neurotransmitters, metabolites, and protons (H^+^), the latter of which activate 3 receptors, GPR4, GPR65, and GPR68 in response to physiologic acidosis (2, 3). Outside of these three limited examples, the effect of pH on GPCR signaling and biology remains largely understudied.

Coincident signals are a well-recognized feature of neuronal and cellular communication (4, 5). By detecting simultaneous inputs, cells can filter signal from noise in complex chemical microenvironments to mount proper responses. As such, the combinatorial, spatial, and temporal nature of coincidence detection enables cells, tissues, and organs to integrate physiologic signals in a manner analogous to sensor array processing. At the molecular level, these activities are mediated by proteins such as NMDA receptor channels, which are synaptic coincidence detectors gated by glutamate and Mg^2+^ inputs that help choreograph the formation of long-term memories (5). Despite their abundance, similar roles for GPCRs as molecular coincidence detectors remain understudied and await exploration. Given that many GPCRs are already known to detect multiple orthosteric and allosteric inputs, there is likely an enormous potential for the discovery, understanding, and therapeutic utilization of molecular coincidence detection within the human GPCRome.

Several lines of evidence suggest that coincidence detection is hard-wired into much of GPCR signaling biology. Recently, we have shown that acid sensing GPCRs GPR4, GPR65, and GPR68 are molecular coincidence detectors of H^+^ and Na^+^ ions. In this case, H^+^ is an activator and Na^+^ is a negative allosteric modulator of GPR4, GPR65, and GPR68 signaling (6). Additionally, GPR68 has recently been shown to integrate H^+^ and mechanical inputs in response to simultaneous extracellular acidification and membrane stretch (7). Other examples include the self-activation of the adhesion receptor, GPR56, that is mediated by coincident collagen-binding and shear force inputs (8), and input-specific synapse formation guided by the latrophilin-3 receptors and its coincident interactions with fibronectin leucine-rich repeat transmembrane proteins and teneurins (9). Here, we show that many other human GPCRs function as Boolean-gated coincidence detectors that couple H^+^ binding to agonism, antagonism, and constitutive activity.

## Results and Discussion

### Yeast are an ideal model system for pH studies of human GPCRs

As transmembrane receptors, GPCRs can simultaneously experience different pH environments established across lipid bilayers. Localizing pH effects to the GPCR-ligand interface thus requires a model system with static intracellular pH (pH_i_) independent of dynamic extracellular pH (pH_e_). Such a model would avoid conflating pH_e_ effects on GPCR signaling with changes in pH_i_ or pH-induced stress responses that affect processes such as intracellular signal transduction and cellular homeostasis. For example, we have previously shown that changes in pH_i_ affect signaling by Gα subunits, the principal transducers of GPCR signals, and small Ras-like GTPases (10, 11). To circumvent such complications and ensure the interpretability of our results, we sought to identify a suitable cell model for studying H^+^ gating of GPCR signaling.

We began by testing the pH tolerance of human embryonic kidney cells (HEK293), the *in vitro* model most often used for cell-based GPCR assays. As shown in Fig. 1A, changes in pH_e_ caused dramatic and unintended pH_i_ alterations in HEK293 cells as reported by the pH biosensor pHluorin (12). This pH intolerance exhibited a sigmoidal shape centered at pH 6.25 and asymptotic pH_i_ values that differed by almost 1 pH unit between high and low pH_e_. In stark contrast, we found that pH_i_ in *S. cerevisiae*, a yeast cell model used for studying human GPCRs (13–15), was independent of pH_e_ (Fig. 1B). Based on this breakthrough, we concluded that the cross-species compatibility and pH tolerance of yeast was ideal for pH studies of human GPCRs. Having already developed a high throughput platform for studying human GPCRs in yeast (15, 16), this new insight provided us with an experimental framework for modeling biological scenarios in which only the GPCR-ligand interface is exposed to large pH changes, such as those that occur in acidified endosomes and tumor microenvironments.

**Fig. 1.**
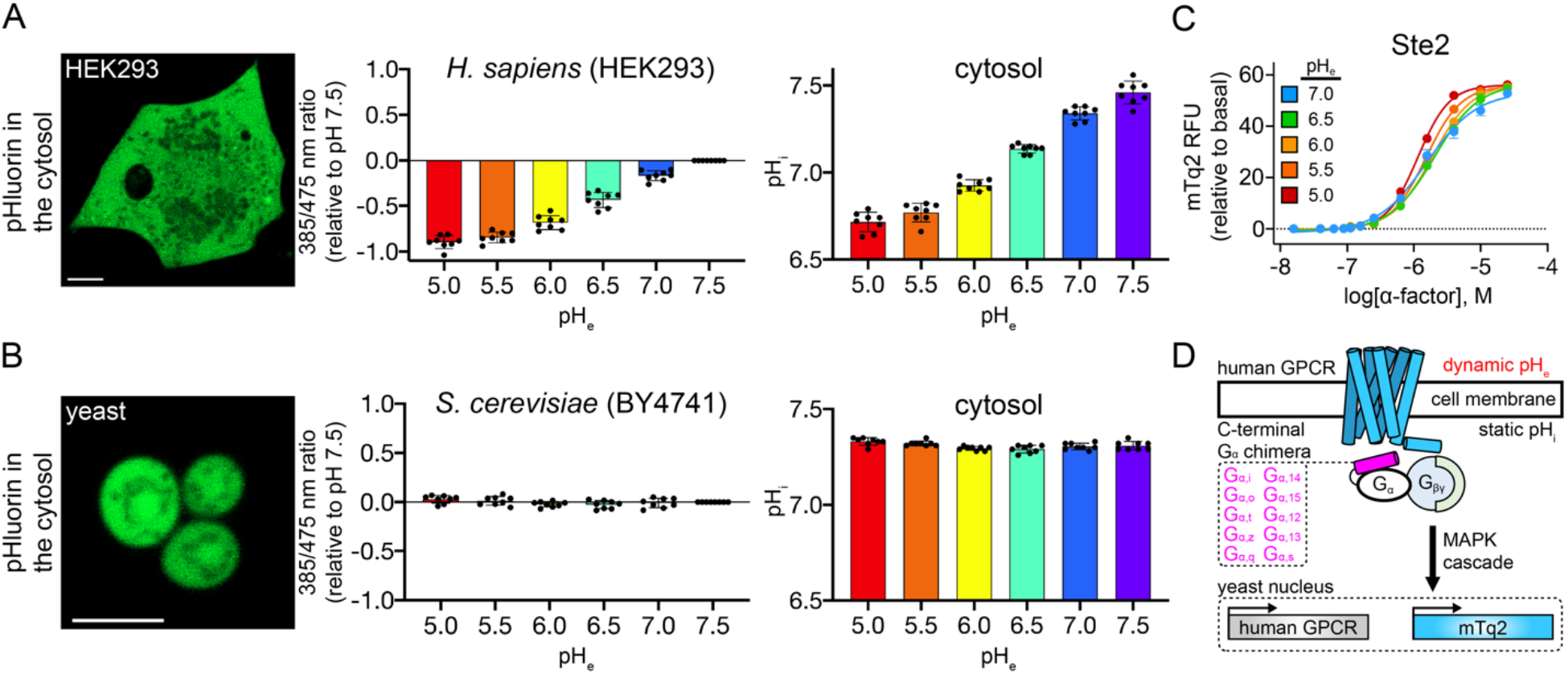
Using yeast to study H^+^-gated coincidence detection by human GPCRs. Confocal images, wavelength ratios, and pH_i_ values reported by the pH biosensor pHluorin in human embryonic kidney (HEK293) (A) and yeast cells (*S. cerevisiae* strain BY4741) (B). Error bars represent SD of n=8 biological replicates. Scale bars, 5 μm. Calibration experiments are provided in Fig. S1. (C) The native yeast GPCR, Ste2, and its downstream pathway components are not affected by changes in pH_e_. Error bars represent SD of n=4 experimental replicates. (D) In the DCyFIR platform, human GPCRs are integrated into a CRISPR-addressable expression cassette in 10 DCyFIR strains, each of which contains a single, unique C-terminal Gα chimera (pink) to test all possible GPCR-Gα coupling combinations. Receptor activation drives the expression of the mTq2 transcriptional reporter protein.

### pH profiling of human GPCRs using the yeast-based DCyFIR platform

The fundamentals of GPCR signaling are conserved from yeast to humans. In mammalian cells, many GPCRs transduce signals through complex networks of intracellular signaling pathways. In contrast, yeast have only one GPCR pathway, controlled by the native yeast receptor Ste2, that is largely insulated from signal crosstalk (17, 18). Unlike human cells, which transduce most GPCR signals through G_α_ subunits, Ste2 signals through the G_βγ_ heterodimer to activate a downstream MAP kinase cascade and transcriptional response. As illustrated in Fig. 1C, we have shown these processes are collectively insensitive to pH_e_. As such, pH responses for human GPCRs in the yeast model cannot be attributed to pH-induced changes in the underlying Ste2 pathway.

As shown in Fig. 1D, we have built a high throughput platform for studying human GPCRs that utilizes the isolated Ste2 pathway: Dynamic Cyan Induction by Functional Integrated Receptors, DCyFIR (6, 15, 16, 19). In the DCyFIR model, activation of the yeast MAP kinase cascade is coupled to a human GPCR by a human/yeast C-terminal Gα chimera (Fig. 1D). To cover all possible human GPCR-Gα combinations, each GPCR is installed into a genome integrated cassette and expressed in 10 strains, each containing a mTurquoise2 (mTq2) transcriptional reporter and unique Gα chimera. Using this platform, we performed the foremost assessment of H^+^-gated GPCR activity by profiling the pH responses of 280 individual GPCR-Gα combinations, covering 28 unique receptors.

### Boolean H^+^ gating of GPCR agonism

As shown in Fig. 2A-B, we screened the set of 280 DCyFIR strains at pH 7 and 5, identifying 131 GPCR-Gα agonist responses. 63 of these individual GPCR-Gα agonist responses exhibited signaling that was exclusive to pH 7 or 5. For example, the somatostatin receptor 5 (SSTR5) and all three sphingosine-1-phosphate receptors (S1PR1, S1PR2, and S1PR3) signaled only at pH 7. We characterized this all-or-nothing H^+^ gating as switch-like Boolean behavior (Fig. 2A). In contrast, the remaining 68 individual GPCR-Gα agonist responses presented in Fig. 2B exhibited non-Boolean behavior by signaling to some degree at both pH 7 and 5. In some cases, such as FFAR2 and LPAR4, agonist responses were more challenging to quantify due to high levels of constitutive activity. In these cases, constitutive activity must be subtracted to observe the Boolean/non-Boolean agonist responses (see Supplemental Fig. S2B). Considering both Boolean and non-Boolean sets, we found that 66% of agonist responses were greater at pH 7, 18% were greater at pH 5, and 16% were similar at both pH values (Fig. 2A-B). Having selected agonists with p*K*_a_ values outside of pH 7 and 5 (Supplemental Dataset 2), these effects were unlikely to be caused by changes in agonist ionization state. These findings show that for many GPCRs, signaling is more likely to be greater at higher pH and dramatically diminished or switched off at lower pH values. However, in rare cases we did find exceptions to this trend (FFAR2 and LPAR4 in Fig. 2A-B).

**Fig. 2.**
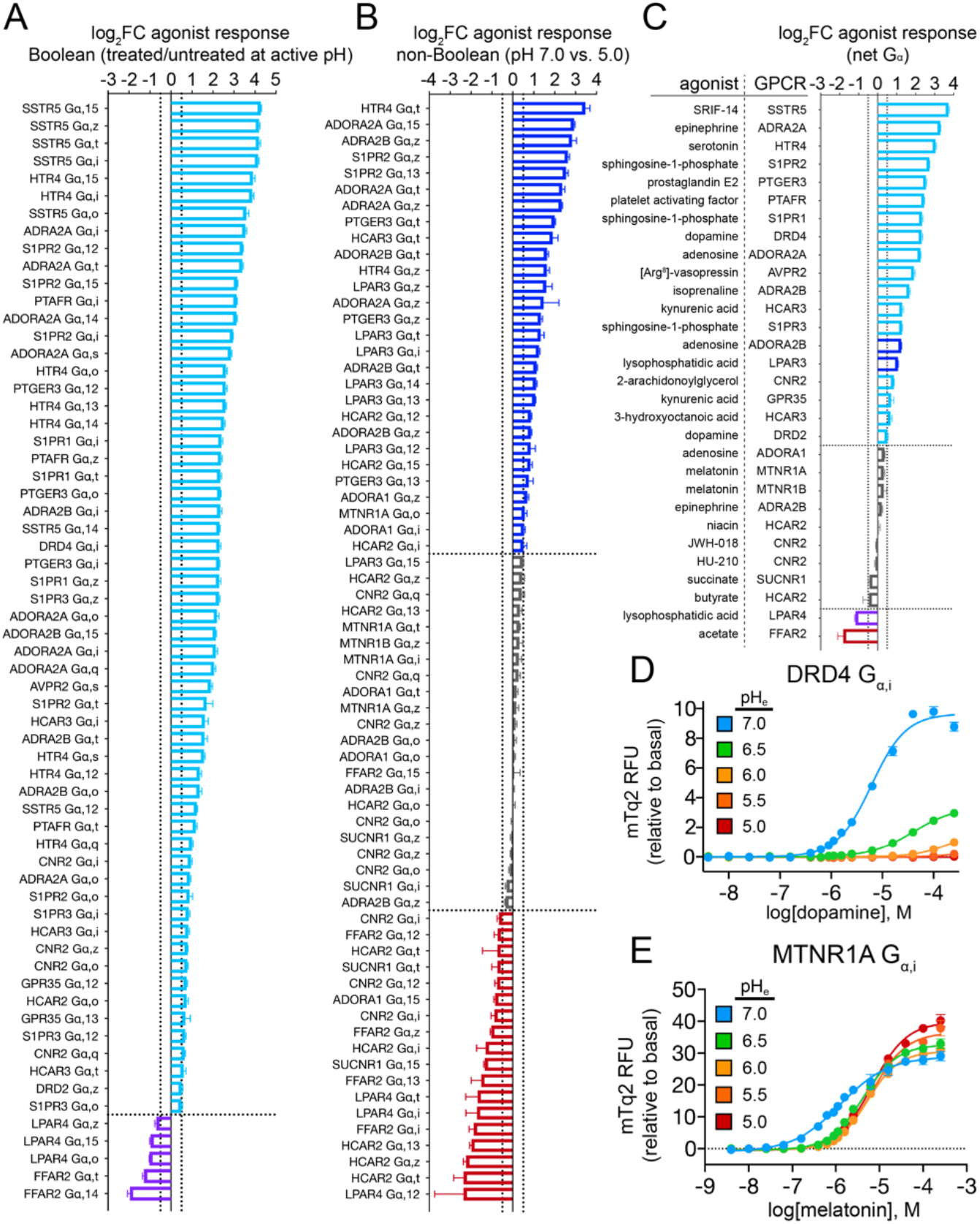
H^+^-gated coincidence detection for 131 GPCR-Gα coupling combinations. Waterfall plots showing Boolean (A) and non-Boolean (B) H^+^-gated agonist responses. Differential agonism was quantified as the log_2_ fold change (log_2_FC) of the ratio of agonist treated and untreated samples at active pH (A) or agonist response between pH 7 and 5 (B). (C) Waterfall plot showing the net Gα agonist responses for a given receptor using the ratio of summed activity for all agonist-active GPCR-Gα strains between pH 7 and 5. Switch-like Boolean responders to high and low pH are colored cyan and purple, respectively. (D) Switch-like Boolean behavior exhibited by dopamine receptor 4 (DRD4 G_α,i_). (E) pH-insensitive GPCR signaling by melatonin receptor 1A (MTNR1A G_α,i_). (A-C) Vertical dashed lines correspond to a log_2_FC = 0.5 and horizontal dashed lines separate examples that signaled more (log_2_FC ≥ 0.5) at pH 7 (blue/cyan bars), similarly (log_2_FC between ± 0.5) between both pH values (gray bars), or more (log_2_FC ≤ 0.5) at pH 5 (red/purple bars). (A, B) Primary screening data is provided in Fig. S2. (A-E) Error bars represent SD of n=4 experimental replicates.

An advantage of the yeast system is that it enabled us to measure the pH responses of individual GPCR-Gα interactions. In most cases, we observed that Gα coupling responses were similar for a given receptor and agonist. For example, all serotonin receptor 4 (HTR4) and lysophosphatidic acid receptor 4 (LPAR4) Gα pairs were more active at pH 7 and 5, respectively. However, Gα coupling patterns were unexpectedly complex in several cases, such as niacin and butyrate agonism of hydroxycarboxylic acid receptor 2 (HCAR2) (Fig. 2A-B and S2). Whereas G_i/o_ coupling was generally stronger at low pH for niacin, G_q_ and G_12/13_ coupling favored high pH. In contrast, butyrate agonism through G_i/o_ and G_13_ generally favored low pH. These observations suggest that H^+^ gating can be allosterically modulated by Gα coupling and that the net effect of H^+^ gating on GPCR agonism could depend on the Gα repertoire expressed in a given cell type.

Our DCyFIR profiling experiments demonstrated that many GPCRs are Boolean and non-Boolean coincidence detectors of H^+^ and agonists. As presented in Fig. 2A-C, this interpretation is consistent for both individual GPCR-Gα pairs and net Gα agonist responses for a given GPCR. As shown in Fig. 2C, the net Gα responses for 18 GPCR-agonist pairs exhibited Boolean behavior, with 17/1 only signaling at pH 7/5, respectively. A striking example of this switch-like H^+^-gated agonism is illustrated by titrations of dopamine receptor 4 (DRD4), for which signaling at pH 7 was switched off at pH 6 (Fig. 2D). Notably, of the 19 net Gα responses more active at pH 7, only 2 exhibited non-Boolean behavior (ADORA2B and LPAR3) (Fig. 2C). In contrast, of the 2 net Gα responses more active at pH 5 (LPAR4 and FFAR2), only LPAR4 exhibited switch-like Boolean signaling. The 9 remaining net Gα responses in Fig. 2C were insensitive to pH, as illustrated in Fig. 2E by titrations of the melatonin receptor 1A (MTNR1A) as a function of pH. Together, these findings suggest that many GPCRs can respond to physiological pH changes like Boolean switches that are turned on and off by high and low H^+^ concentrations.

### All modes of GPCR pharmacology can be regulated by H^+^-gated coincidence detection

As shown in Fig. 3, we next investigated the effects of H^+^ gating on the major modes of GPCR pharmacology. Using a set of representative receptors and select Gα strains, we demonstrated that H^+^ inputs can modulate agonist potency and efficacy (Fig. 3A-D), and receptor sensitivity and inhibition (Fig. 3E and 3F, respectively). As shown in Fig. 3A, the α_2B_-adrenergic receptor (ADRA2B) appears to be insensitive to pH at high concentrations of the agonist epinephrine. However, pH has a profound effect on ADRA2B agonism at lower epinephrine concentrations, with increasingly lower pH values reducing epinephrine potency (pEC_50_ values) by >2 orders of magnitude (Fig. 3A). A consequence of these pH dependent effects on epinephrine potency is that H^+^-gated agonism of ADRA2B exhibited near-Boolean behavior between pH 7 and 5, however only at concentrations between ~10 and ~1000 nanomolar. This finding illustrates that H^+^ modulation of GPCR activity, or apparent lack thereof, can depend on agonist concentration.

**Fig. 3.**
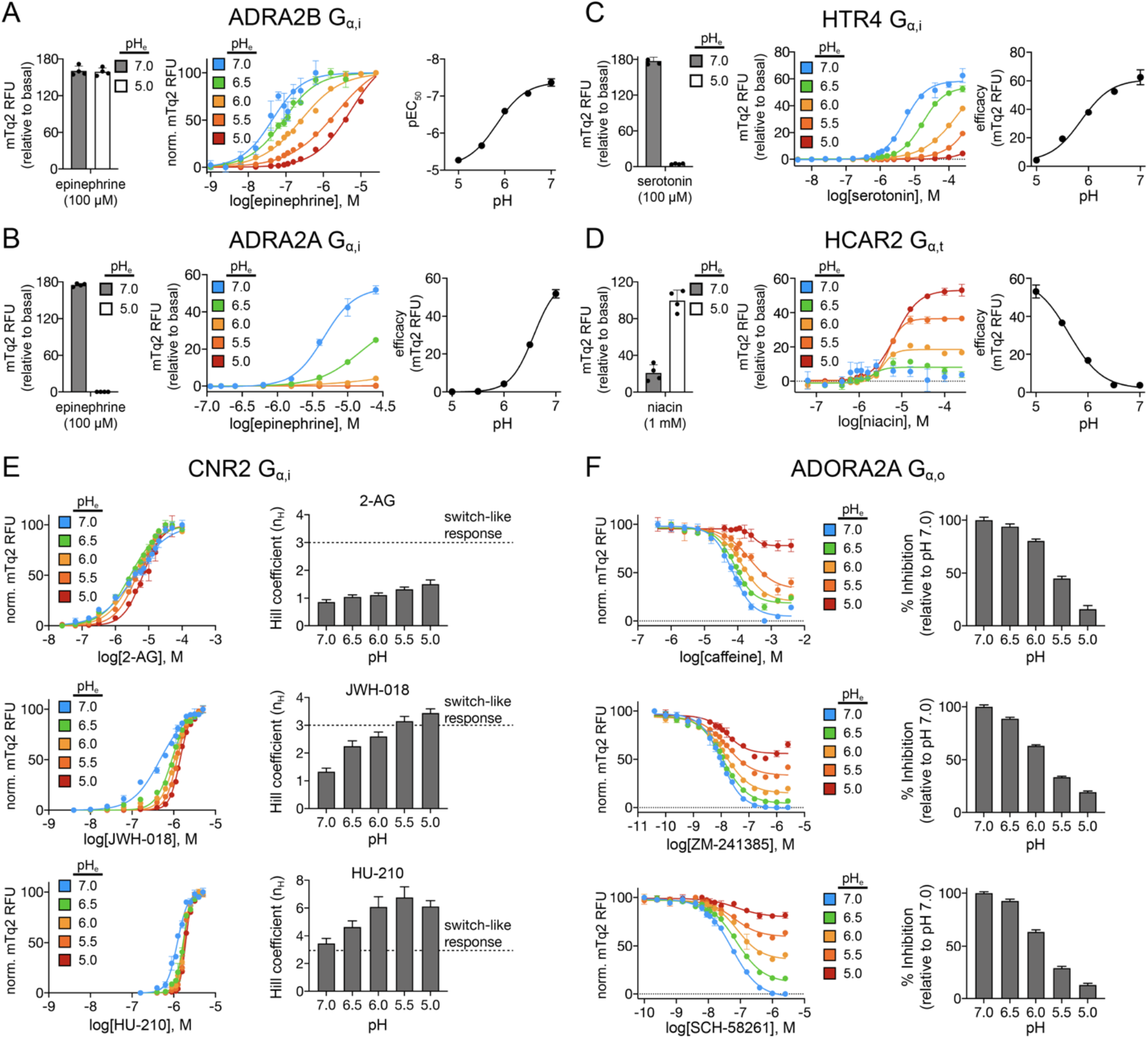
Pharmacological modalities of H^+^-gated coincidence detection by GPCRs. (A) Potentiation of agonist potency. Endpoint measurements of ADRA2B agonism (left) at pH 7 (gray bars) and 5 (white bars). Epinephrine dose-response curves (middle) and pEC_50_ values (right) for ADRA2B. (B-D) Potentiation of agonist efficacy. Endpoint measurements (left) of ADRA2A (B), HTR4 (C), and HCAR2 (D) agonism at pH 7 (gray bars) and 5 (white bars). Dose-response curves (middle) and efficacy (right) of each GPCR with its endogenous agonist. (E) Potentiation of sensitivity. 2-arachidonoyl glycerol (2-AG) (top), JWH-018 (middle), and HU-210 (bottom) dose-response curves (left) and Hill coefficients (right) for CNR2. (F) Potentiation of inhibition/antagonism. Caffeine (top), ZM-241385 (middle), and SCH-58261 (bottom) dose-response curves (left) and percent inhibition (right) for ADORA2A in the presence of 10 μM adenosine. (A-F) Error bars represent SD of n=4 experimental replicates.

In the case of ADRA2B, epinephrine elicits full agonist responses between pH 7 and 5 (Fig. 3A). As such, the set of ADRA2B dose-response curves all achieved full terminal efficacy at high epinephrine concentration. In contrast, epinephrine agonism of the α_2A_-adrenergic receptor (ADRA2A) was weaker and only reached full terminal efficacy at pH 7 (Fig. 3B). At lower pH, ADRA2A achieved what we refer to as apparent terminal efficacy (at pH 6.5) or did not signal. As a result, H^+^-gated ADRA2A agonism exhibited switch-like Boolean behavior between pH 7 and 6 at micromolar epinephrine concentrations. The concept of terminal efficacy versus apparent terminal efficacy is further illustrated in Fig. 3C for serotonin receptor 4 (HTR4). In this case, serotonin agonism of HTR4 achieved terminal efficacy at pH 6.5 and above, and apparent terminal efficacy at pH 6.0 and below, leading to switch-like Boolean H^+^ gating at micromolar serotonin concentrations. Lastly, as shown for hydroxycarboxylic acid receptor 2 (HCAR2), the effect of pH on agonism can be strictly limited to terminal efficacy (Fig. 3D). In this case, niacin agonism of HCAR2 maintained the same pEC_50_ value at all pH values yet displayed a dramatic increase in HCAR2 terminal efficacy with decreasing pH.

As illustrated in Fig. 3E-F, H^+^-gated coincidence detection also extends to GPCR sensitization and inhibition. Fig. 3E shows how pH modulates cannabinoid receptor 2 (CNR2) responses to endogenous (2-arachidonoylglycerol, 2-AG) and synthetic (JWH-018 and HU-210) agonists. In the case of 2-AG, sensitivity of CNR2 signaling was non-cooperative and unaffected by pH as indicated by a near constant Hill coefficient (*nH*) between 0.9 to 1.5. In contrast, H^+^ gating of JWH-018 and HU-210 agonism elicited increasingly switch-like ultrasensitive responses (*n*_H_ > 2) (20) as pH was decreased. As shown in Fig. 3F, we observed a similar trend in H^+^ gating of adenosine receptor A2A (ADORA2A) antagonism using the known competitive inhibitors caffeine, ZM-241385, and SCH-58261 in the presence of 10 μM adenosine. All three inhibitors were 50% effective at pH 6 and were almost completely ineffective at pH 5.0. Based on these findings, we concluded that the potential for H^+^ gating of GPCR pharmacology warrants far more attention in the development and evaluation of new and existing therapeutics. Our findings further suggest that as a principle of drug design, intentional H^+^ gating may find broad utility in the development of drugs that target GPCRs exclusively in acidic microenvironments such as endosomes and tumors, and vice versa.

### New GPCR proton sensors

As shown by the waterfall plots in Fig. 4A-B, 48 of the 280 individual GPCR-Gα strains screened at pH 7 and 5 were constitutively active, with 24 responses showing switch-like Boolean behavior at pH 7 or 5 (Fig. 4A). For example, the known acid sensor GPR68 signaled exclusively below pH 7. The 24 remaining GPCR-Gα responses in Fig. 4B exhibited non-Boolean behavior by signaling to some degree at both pH 7 and 5. Considering both Boolean and non-Boolean sets, we found that 33% signaled higher at pH 7, 54% signaled higher at pH 5, and 13% were similar at both pH values (Fig. 4A-B). Unlike H^+^ gating of GPCR agonism (Fig. 2), these findings show that GPCR constitutive activity is more likely to be greater at low pH in our receptor set. This result can be attributed to the high degree of Gα-coupling promiscuity we observed for known acid-sensing GPCRs (Fig. 4A-B, solid bars).

**Fig. 4.**
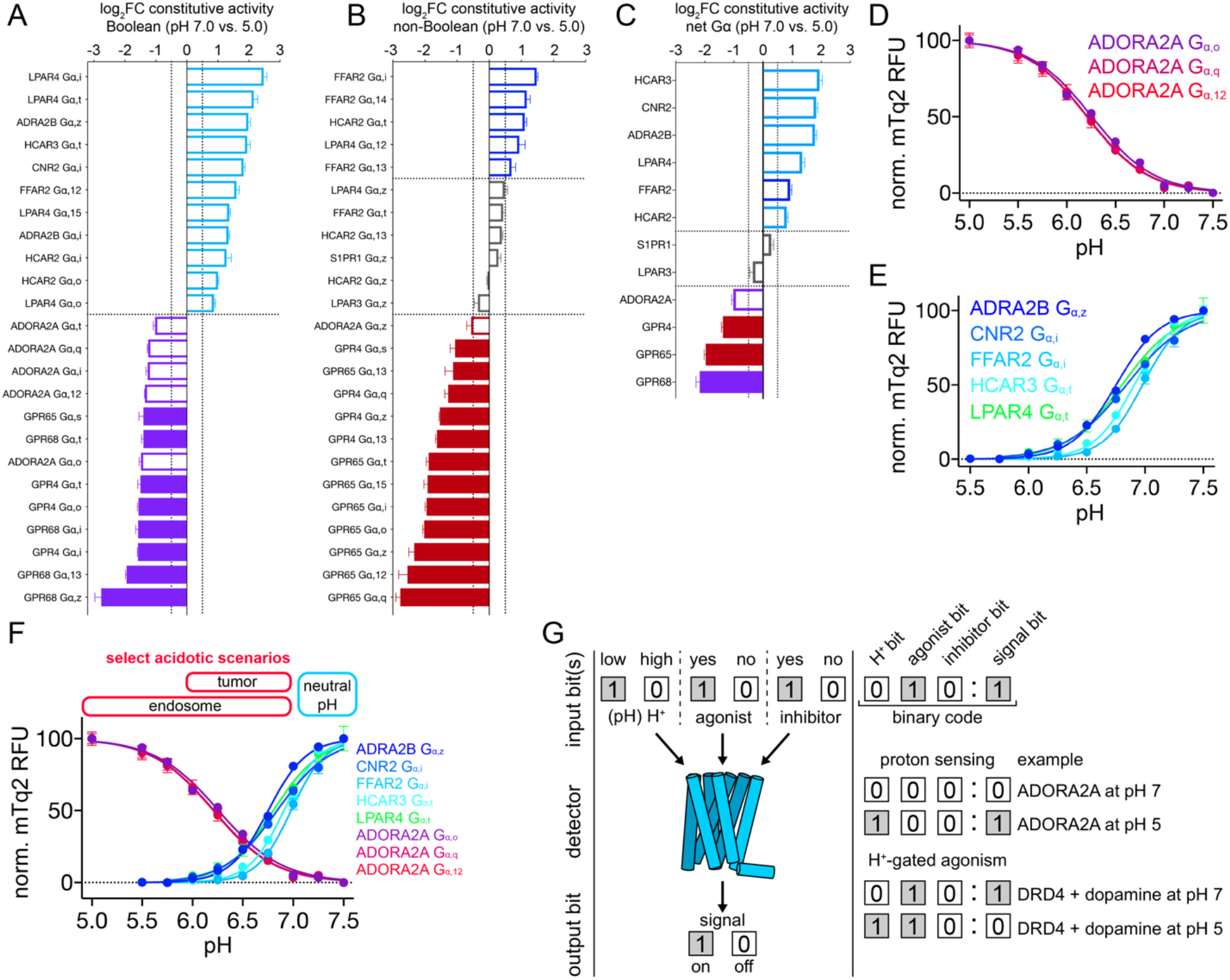
New pH sensors and Boolean logic of H^+^-gated GPCR signaling. (A, B) Waterfall plots showing Boolean (A) and non-Boolean (B) instances of H^+^-gated GPCR-Gα constitutive activity. Differential signaling was quantified as the log_2_FC between pH 7 and 5. (C) Waterfall plot showing the net Gα responses for a given receptor using the ratio of summed mTq2 RFU for all constitutively active GPCR-Gα strains between pH 7 and 5. Switch-like Boolean responders to high and low pH are colored cyan and purple, respectively. (D-E) New proton sensors activated by low (D) and high (E) pH. (F) pH titrations of new proton-sensing GPCRs in the context of pH changes associated with select physiological processes and pathologies. (G) An illustration using ADORA2A proton sensing and H^+^ gating of DRD4 agonism to show how pH effects on GPCRs can be described by a Boolean language expressed in binary code. (A-C) Vertical dashed lines correspond to a log_2_FC = 0.5 and horizontal dashed lines separate examples that signaled more (log_2_FC ≥ 0.5) at pH 7 (blue/cyan bars), similarly (log_2_FC between ± 0.5) between both pH values (gray bars), or more (log_2_FC ≤ 0.5) at pH 5 (red/purple bars). Filled bars correspond to strains of the known acid sensors (GPR4, GPR65, and GPR68). (A-F) Error bars represent SD of n=4 experimental replicates.

As shown in Fig. 4C, net Gα signaling for 7 of 12 constitutively active GPCRs exhibited Boolean-like behavior. Of these 7 receptors, 5 were activated by high pH (HCAR3, CNR2, ADRA2B, LPAR4, and HCAR2) and 2 by low pH (GPR68 and ADORA2A). Only 1 of the 12 receptors (FFAR2) exhibited non-Boolean behavior and higher signaling at pH 7 (Fig. 4C). Notably, H^+^ gating of LPAR4 and FFAR2 constitutive activity (greater at pH 7) was the inverse of their net Gα agonist responses (greater at pH 5) shown in Fig. 2C. This finding illuminates previously unknown complexities involving GPCR signaling and pH that likely extend to many other receptors. As shown by the pH titrations in Fig. 4D-E, we discovered a new acid sensor, ADORA2A (Fig. 4D), and a new class of H^+^ sensors that are activated by high pH (Fig. 4E).

### The Boolean-like language of proton sensing and H^+^-gated GPCR coincidence detection

Our findings demonstrate that proton sensing and H^+^-gated agonism are recurring features of GPCR signaling biology. Remarkably, the midpoint of proton-sensing and H^+^-gated responses (pH_50_ values) are tightly grouped within the physiologic pH range, having an average value of 6.25 ± 0.75 pH units (collectively for the pH titrations in Figs. 2-4). Focusing on the newly discovered acid sensor, ADORA2A, pH titrations through three GPCR-Gα pairs are in near-perfect agreement, all having pH_50_ values of 6.20 ± 0.03. This is similar to the pH_50_ of 6.35 ± 0.02 for the known acid sensor GPR68, which we have studied extensively (6). As shown in Fig. 4F, acidotic biological scenarios should activate ADORA2A in the absence of adenosine, such as situations in which ADORA2A is constitutively endocytosed (21). In contrast, our findings suggest that acidotic scenarios should negatively modulate most GPCR agonism (Fig. 2). For example, the switch-like Boolean inactivation of DRD4 between low and high pH could be a factor in the regulation of dopamine receptor signaling from endosomes (22, 23).

In summary, we have shown that much of GPCR proton sensing and H^+^-gated coincidence detection exhibits switch-like Boolean characteristics in the physiologic pH range. This concept is illustrated in Fig. 4G, in which we show that the effects of pH on many GPCRs can be described by a Boolean language expressed in concise binary codes. Moving forward, this formalism can be expanded to include a variety of additional inputs, such as allosteric modulators, to classify the complexities of GPCR logic gating. As such, we believe the technical and conceptual advances embodied in this work establish a new paradigm for understanding and exploring signaling in acidified microenvironments such as endosomes, tumors, synapses, and ischemic vasculature that will likely extend well beyond the GPCR superfamily.

## Materials and Methods

### Media

All yeast strains were struck to YPD media plates (20 g/L peptone, 10 g/L yeast extract, 2% glucose, 15 g/L agar) from 30% glycerol stock cultures stored at −80 °C. For all DCyFIRscreen experiments and pH titrations, yeast were grown in low-fluorescence synthetic complete dropout media (SCD complete media; 50 mM potassium phosphate dibasic, 50 mM MES hydrate, 5 g/L ammonium sulfate, 1.7 g/L yeast nitrogen base without amino acids, folic acid, and riboflavin (Formedium; CYN6505), 0.79 g/L complete amino acid mix (MP Biomedicals; 4500022), and 2% glucose) titrated to the desired pH using HCl or NaOH and filter-sterilized. For yeast work using pHluorin to measure intracellular pH, yeast were selected for on plates lacking leucine and grown in SCD media lacking leucine (5 g/L ammonium sulfate, 1.7 g/L yeast nitrogen base without amino acids (MP Biomedicals; 4510522), 1 NaOH pellet (VWR; BDH9292), 0.69 g/L CSM-LEU (amino acid mix lacking leucine), 2% glucose, and 15 g/L Bacto Agar for plates) that was filter-sterilized. For pHluorin purification experiments, autoinduction media was prepared by combining 800 mL ZY media (10 g/L tryptone, 5 g/L yeast extract), 16 mL 50Χ 5052 (25% (w/v) glycerol, 2.5% (w/v) glucose, 10 % (w/v) α-lactose), 16 mL 50Χ M (1.25 M Na_2_HPO_4_, 1.25 M KH_2_PO_4_, 2.5 M NH_4_Cl, 0.25 M Na_2_SO_4_), and 1.6 mL 1 M MgSO_4_. Phosphate buffered saline (PBS) was made by first preparing solutions of mono- and dibasic potassium phosphate (100 mM potassium chloride, 25 mM potassium phosphate). Individual pH solutions of PBS (pH = 5.0-8.5, increments of 0.25) were prepared by mixing the mono- and dibasic solutions until the desired pH was reached, measured using an Accumet XL150 pH meter (Fisher Scientific; Hampton, NH). PBS-TCEP was prepared as PBS (pH = 7.0) with 1 mM TCEP, 0.15 mM PMSF, 1 mM MgCl_2_, 1 mM EDTA, 0.1% Triton, spiked with lysozyme and 10 μL DNAse.

### Plasmids

Yeast plasmid pYEplac181-pHluorin was a gift from the Rajini Rao lab at Johns Hopkins. The pHluorin gene from pYEplac181-pHluorin was subcloned into a pLIC-His vector (pLIC-His-pHluorin) for recombinant overexpression and purification from *E. coli*. Mammalian plasmid pcDNA3.1(+)-pHluorin was made by synthesizing and cloning a codon optimized sequence of pHluorin into the pcDNA3.1(+) backbone (GenScript; Piscataway, NJ).

### Strains/Cell lines

#### Yeast

Yeast strains used in this work were previously described in another study (15). Briefly, wild-type BY4741 was modified using CRISPR/Cas9 to prime the endogenous pheromone pathway (*far1∆, sst2∆, ste2∆*) for expression of human GPCRs directly from the yeast genome (installation of an expression cassette (*PTEF*, *TCYC1B*) on chromosome X), enable coupling of the yeast Gα to human GPCRs (via chimeric Gα proteins encoding the five C-terminal amino acids from human Gα proteins), and link receptor activation to a transcriptional reporter, mTurquoise2 (*fig1∆::mTurquoise2*). For a comprehensive list of all strains, see Dataset S1.

#### Mammalian

HEK293T (HEK293T/17) cells were purchased from American Type Culture Collections (ATCC; Gaithersburg, MD) and maintained in DMEM (Thermo Fisher; 11995-065) supplemented with 10% Fetal Bovine Serum (Thermo Fisher; 10082-147) and 1% Penicillin-Streptomycin (Thermo Fisher; 15140-122).

### Intracellular pH measurements

#### pHluorin purification

Ratiometric pHluorin (12) was overexpressed in E. coli using autoinduction. First, 25 ng pLIC-His-pHluorin was transformed into competent BL21(DE3) RIPL cells. Transformants were grown overnight in 5 mL LB + carbenicillin (100 μg/mL; Sigma-Aldrich; C1389). The next morning, the overnight culture was transferred into 800 mL autoinduction media (see “*Media*” above), grown at 37 °C with shaking (200 rpm) for 8 hours, then grown overnight at 18 °C with shaking (200 rpm). The 800 mL autoinduction culture was split over two 500 mL centrifuge bottles and cells were harvested by centrifugation at 4500 rpm for 30 minutes. The supernatant was removed, and the pellet was resuspended in 35 mL PBS, transferred to 50 mL tubes, and stored at −20 °C. To purify overexpressed ratiometric pHluorin, cell pellets were thawed on ice and treated with TCEP, MgCl_2_, EDTA (all to 1 mM), PMSF (to 0.15 mM), Triton (to 0.1%), lysozyme, and 10 μL DNAse, then lysed using a NanoDeBEE homogenizer (BEE International; South Easton, Massachusetts) at 30,000 psi. Cell lysate was transferred to a 50 mL tube and centrifuged at 15,000 rpm for 30 minutes. Supernatant was then transferred to a 15 mL tube and incubated with 1 mL His bead slurry (Sigma-Aldrich; P6611) for 20 minutes at 4 °C. Beads were harvested by centrifugation at 1000 rpm for 1 minute, then washed five times with PBS-TCEP (see “*Media*” above). Beads were then transferred to a 2 mL spin column (Thermo Scientific; 89896) and eluted with PBS-TCEP + 200 mM imidazole. Eluted protein was collected in a 15 mL tube by centrifugation at 1000 rpm for 1 minute, transferred to a slide-a-lyzer dialysis cassette (Thermo Scientific; 87730), and dialyzed overnight in 4 L PBS-TCEP at 4 °C. Purified pHluorin stocks were stored at −80 °C.

#### pHluorin calibration

A pHluorin standard curve was calculated using purified ratiometric pHluorin. Briefly, 2 μL of ratiometric pHluorin (100 μM) was resuspended in 198 μL PBS buffer titrated to each pH (see “*Media”* above) and 50 μL aliquots were moved into a 96-well microplate (CytoOne; CC7626-7596) in technical triplicate. Excitation spectra were collected using a ClarioStar microplate reader (BMG LabTech; Offenburg, Germany) with the following parameters: (top read, 40 flashes/well, excitation start: 340-10 nm, excitation end: 495-10 nm, 1 nm steps; emission: 520-10 nm; instrument gain: 1700). Raw fluorescence values at 385 and 475 nm were used to calculate the pHluorin ratio (385/475 nm) for each replicate. A standard curve was built by plotting the pHluorin ratio as a function of pH. Data was fit (sigmoidal, 4PL, X is log[concentration]) using GraphPad Prism (San Diego, California).

#### Measuring yeast intracellular pH

Yeast intracellular pH measurements were collected using the pH biosensor ratiometric pHluorin expressed in the cytosol. 150 ng of plasmid pYEplac181-pHluorin was transformed into the wild-type yeast BY4741 using standard methods. Transformed cells were selected for on SD plates lacking leucine (see “*Media*” above). Eight individual colonies expressing cytosolic pHluorin were grown to mid-log phase in SCD media lacking leucine. Cells were harvested and washed three times in SCD Complete Media titrated to various pH values (5.0, 5.5, 6.0, 6.5, 7.0, 7.5, 8.0), then transferred to a 384-well plate (Greiner; 781096). Spectral scans were collected using a ClarioStar microplate reader with the following parameters: top read, 40 flashes/well, excitation start: 340-10 nm, excitation end: 495-10 nm, 5 nm steps; dichroic: 430 nm; emission filter: 520-10 nm. pHluorin spectral scans were generated by subtracting the background autofluorescence of media at each pH. pHluorin ratios were calculated from pHluorin spectral scans for each replicate by dividing the fluorescence at 385 nm by the fluorescence at 475 nm. Intracellular pH was quantified using the pHluorin ratios as previously described (10).

#### Measuring mammalian cell intracellular pH

Mammalian cell intracellular pH measurements were collected using the pH biosensor ratiometric pHluorin expressed in the cytosol. Prior to transfection, HEK293T cells were seeded into 6-well plates (CytoOne; CC7682-7506) at a density of 700,000 to 800,000 cells per well. Four hours later, cells were transfected with 2 μg plasmid DNA (pcDNA3.1(+) empty vector; n=1 well, or pcDNA3.1(+)-pHluorin; n=8 wells) complexed with 6 μL TransIT-2020 transfection reagent (Mirus Bio; MIR 5404) at a final DNA concentration of 8 ng/uL in Advanced DMEM (Thermo Fisher; 12491-015). After 72 hours, cells were detached using TrypLE (Thermo Fisher; 12604-013), resuspended in 1 mL media (DMEM; 10% FBS, 1% PenStrep), and transferred to a 1.5 mL tube. Cells were harvested by centrifugation (600 rpm, 3 minutes) and resuspended in 600 μL 1Χ HBSS (20 mM MES, 20 mM HEPES). 100 μL cell suspension was transferred to 6 1.5 mL tubes and harvested by centrifugation (600 rpm, 3 minutes). Cells were washed with 1Χ HBSS (20 mM MES, 20 mM HEPES) titrated to each pH (5.0, 5.5, 6.0, 6.5, 7.0, and 7.5) and transferred into a 384-well plate. A spectral scan of each well was collected using a ClarioStar microplate reader using the settings described above in “*Measuring yeast intracellular pH”*. pHluorin spectral scans were generated by subtracting the background autofluorescence of cells transfected with pcDNA3.1(+) from those transfected with pcDNA3.1(+)-pHluorin. pHluorin ratios were calculated from pHluorin spectral scans for each replicate by dividing the fluorescence at 385 nm by the fluorescence at 475 nm. Intracellular pH was quantified using the pHluorin ratios as previously described (10).

### DCyFIRscreen profiling of GPCR proton-linked coincidence detection

For all DCyFIRscreen experiments in this study, individual strains were grown to mid-log phase in SCD complete media titrated to either pH 5.0 or 7.0. Cells were harvested and normalized to a starting OD_600_ = 0.1 before treatment with vehicle or agonist in both pH medias. Cells were incubated at 30 °C until cell densities between the two pH medias were similar (OD_600_ ~ 3-4; ~18 hours at pH 5.0, ~24-48 hours at pH 7.0) before mTq2 fluorescence was measured using a ClarioStar microplate reader (bottom read, 10 flashes/well, excitation filter: 430–10 nm; dichroic filter: LP 458 nm; emission filter: 482–16 nm; gain = 1,300). mTq2 fluorescence for treated/untreated cells at pH 5.0 and 7.0 was plotted using GraphPad Prism. For plots displaying DCyFIRscreen data “relative to basal”, mTq2 fluorescence from untreated cells was used as baseline and subtracted from the mTq2 fluorescence from treated cells.

### Analysis of raw DCyFIRscreen data

#### Agonist response

In total, 340 DCyFIRscreen experiments covering 28 unique receptors (280 GPCR-Gα strains) treated with vehicle or agonist(s) were analyzed. A given GPCR-Gα strain was considered responsive to agonist if treated cells exhibited a minimal mTq2 fluorescence ≥ 25,000 RFU and if the log_2_ fold-change (log_2_FC) of mTq2 RFU from treated relative to untreated cells was ≥ 0.5 at either pH 5.0 or 7.0. In one case (ADORA2A G_α,12_), we observed more mTq2 fluorescence from untreated cells than treated cells. Although this strain met the requirements described above, the ligands used in the screen were strictly agonists. As such, this strain was removed from the set of agonist-responsive GPCR-Gα strains. We also excluded another strain (ADRA2B G_α,13_ treated with epinephrine) from the set of agonist-responsive GPCR-Gα strains after it was experimentally determined to be a false-positive (data not shown). Using these criteria, we identified 131 agonist-responsive GPCR-Gα strains and calculated their raw agonist response (treated mTq2 RFU – untreated mTq2 RFU) for each replicate at pH 7.0 and 5.0.

#### Agonist response, Boolean criteria

An agonist-responsive GPCR-Gα strain was considered to exhibit Boolean-like behavior if the mean raw agonist response at one pH was near zero (e.g. less than the mean of the corresponding vehicle-treated controls), suggesting all-or-nothing agonism between both pH values. In a small subset of cases where agonist response was weak and appeared to be Boolean (ADORA1 G_α,15_ + adenosine, CNR2 G_α,12_ + HU-210, PTGER3 G_α,13_+ prostaglandin E2, LPAR3 G_α,12_ + LPA, and FFAR2 G_α,15_+ acetate), we could not rule out that a lack of signaling at either pH was due to weak (e.g. all-or-nothing) signaling. In these cases, and consistent with our conservative approach to classifying Boolean-like behavior, we labeled these examples as non-Boolean. Using these criteria, we identified 63 agonist-responsive GPCR-Gα strains that exhibited Boolean-like behavior, and 68 that exhibited non-Boolean-like behavior. Differential agonism was then determined by calculating the log_2_FC of mTq2 RFU from treated cells relative to untreated cells at active pH (for Boolean-like responses) or raw agonist responses between cells treated at pH 7.0 relative to pH 5.0 (for non-Boolean-like responses) for all replicates.

#### Net Gα agonist response

For a given receptor-agonist pair, net Gα agonist response was calculated as the sum of mTq2 RFU from treated and untreated agonist-responsive GPCR-Gα strains for each replicate at pH 5.0 and 7.0. The raw net Gα agonist response was determined by subtracting the net Gα agonist response of untreated cells from treated cells at pH 5.0 and 7.0 for each replicate. A net Gα agonist response was considered to exhibit Boolean-like behavior if the mean raw net Gα agonist response at one pH was near zero (e.g. less than the sum of the corresponding untreated controls). Using these criteria, we identified 18 net Gα agonist responses that exhibited Boolean-like behavior and 12 that exhibited non-Boolean-like behavior. Differential agonism of net Gα agonist response was determined by calculating the log_2_FC of the net Gα agonist response from treated cells relative to untreated cells at active pH (for Boolean-like responses) or from the raw net Gα agonist response between cells treated at pH 7.0 and pH 5.0 (for non-Boolean-like responses) for all replicates.

#### Constitutive activity

In total, 28 unique receptors (280 GPCR-Gα strains) were assessed for constitutive activity. A given GPCR-Gα strain was considered constitutively active if the log_2_FC of the mean of untreated cells relative to untreated controls was ≥ 1.0. In one case (ADORA1 G_α,t_), we excluded a GPCR-Gα strain from the set of constitutively active strains after it was experimentally determined to be a false positive (data not shown). This selection criteria identified 48 unique constitutively active GPCR-Gα strains.

#### Constitutive activity, Boolean criteria

A constitutively active GPCR-Gα strain was considered to exhibit Boolean-like behavior if the mean mTq2 RFU for untreated cells at one pH was near zero (e.g. less than twice the mTq2 RFU of untreated controls). Using these criteria, we identified 24 GPCR-Gα strains that exhibit Boolean-like constitutive activity and 24 that exhibited non-Boolean-like constitutive activity. Differential constitutive activity was quantified by calculating the log_2_FC of the mTq2 RFU between untreated cells at pH 7.0 and 5.0 for each replicate.

#### Net Gα constitutive activity

Net Gα constitutive activity was determined by calculating the sum of mTq2 RFU for all constitutively active GPCR-Gα strains for a given receptor at pH 7.0 and 5.0 for each replicate. The constitutive activity of a given GPCR was considered to exhibit Boolean-like behavior if the mean net Gα constitutive activity at one pH was near zero (e.g. less than twice the mean mTq2 RFU of the corresponding untreated controls). Using these criteria, we identified 7 receptors that exhibited Boolean-like net Gα constitutive activity and 3 that exhibited non-Boolean-like constitutive activity. In the specific case of ADORA2A, 5 of the 6 constitutively active GPCR-Gα strains exhibited Boolean-like responses. Based on this finding, and on our conservative approach for characterizing the effects of pH on receptor behaviors, we classified ADORA2A as Boolean. Differential net Gα constitutive activity was determined by calculating the log_2_FC of the sum of untreated constitutively active GPCR-Gα strains at pH 7.0 relative to 5.0 for each replicate.

### Ligand and constitutive activity titrations

#### Ligand titrations as a function of pH

Individual strains were grown to mid-log phase in SCD complete media titrated to each pH (5.0, 5.5, 6.0, 6.5, and 7.0). Cells were harvested and normalized to a starting OD_600_ = 0.1 in each pH media. 36 μL normalized cells at each pH were moved to a 384-well microplate with 4 μL 10Χ ligand stocks in a 16-point serial dilution series centered around the apparent pEC_50_ at pH = 7.0. Plates were sealed with a permeable lid (Diversified Biotech; BERM-2000) and incubated at 30 °C. At least seven fluorescence and absorbance measurements were recorded in 1-3 hr intervals over ~24 hours. Fluorescence values were plotted as a function of absorbance for each ligand concentration at each pH and fit (simple linear regression) using GraphPad Prism. Slope values from the fit function were then plotted for each ligand concentration at each pH. At least two points from the relative baseline of each curve were averaged and used as baseline for background correction (e.g. “relative to basal”). Data were fit (log(agonist) vs. response -- Variable slope (four parameters)) using GraphPad Prism. In cases where data was normalized from 0-100%, constraints of bottom = 0 and top = 100 were used in the fit function.

#### Constitutive activity titrations as a function of pH

Individual strains were grown to mid-log phase in SCD complete media titrated to pH 7.0 (for strains with higher constitutive activity at low pH) or to pH 5.0 (for strains with higher constitutive activity at high pH). Cells were harvested and normalized to a starting OD_600_ = 0.5 then washed twice in SCD complete media titrated to each pH (5.0, 5.5, 5.75, 6.0, 6.25, 6.5, 6.75, 7.0, 7.25, and 7.5). 40 μL cells were moved into a 384-well microplate, sealed with a permeable lid, and incubated at 30 °C. Fluorescence and absorbance measurements were collected, and data was analyzed as described above in “*Ligand titrations as a function of pH”.*

### Confocal Microscopy

#### Yeast

Wild-type yeast transformed with pYEplac181-pHluorin (see “*Measuring yeast intracellular* pH” above for details) were grown to mid-log phase in media lacking leucine. Cells were normalized to an OD_600_ = 1.0 in low fluorescence media (see “*Media*” above), transferred to an ~1 cm square agar “pad” on a glass microscope slide (VWR; 16004-422) and covered with a glass coverslip (VWR; 48366-227). Fluorescence images were collected using an LSM800 confocal microscope (Zeiss; Oberkochen, Germany) with the following parameters: pinhole 1.00 AU/44 μm; laser wavelength 488 nm, 1.0% intensity; 495 nm excitation; 520 nm emission.

#### HEK293T

Cells transfected with pcDNA3.1(+)-pHluorin (see “*Measuring mammalian intracellular pH*” above for details) were detached with TrypLE (Thermo Fisher; 12604-013) and transferred to confocal dishes (VWR; 75856-742) at a density of 300,000 cells/dish in a total of 3 mL media (DMEM; 10% FBS, 1% PenStrep). After 48 hours, fluorescence images were collected using an LSM800 confocal microscope (Zeiss; Oberkochen, Germany) with the following parameters: pinhole 0.67 AU/32 μm; laser wavelength 488 nm, 3.0% intensity; 495 nm excitation; 520 nm emission.

## Acknowledgements

This work was supported by the National Institutes of Health through the National Institute of General Medical Sciences (R35GM119518) to D.G.I.

## Author Contributions

D.G.I. and N.J.K. wrote the manuscript. N.J.K. engineered yeast strains and performed the experiments. J.B.R., G.J.T., and C.R.O. engineered yeast strains. W.M.M. provided technical support. J.B.R. and G.J.T. reviewed and revised the manuscript.

## Data availability.

All relevant data, protocols, results, and analyses are available in the main text. All computer code and yeast strains associated with this work are available upon request.

