## Supplemental Information for "Proton-gated coincidence detection is a common feature of GPCR signaling"

### Supplementary Information

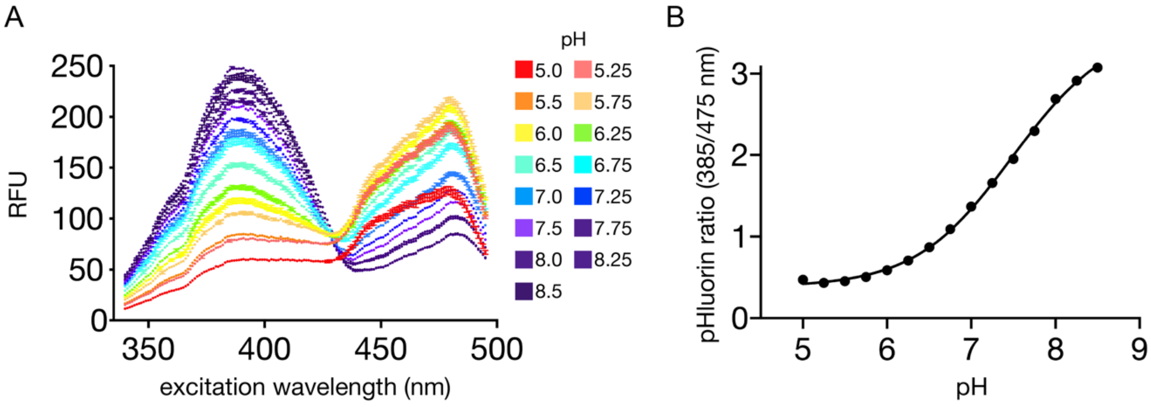


**Figure S1. pHluorin excitation spectrum and standard curve (Related to Fig. 1).**

(A) Excitation spectrum of purified ratiometric pHluorin. Error bars represent SD of n=3 technical replicates.

(B) Standard curve calculated by the fluorescence ratio (ex: 385/475 nm, em: 520 nm) of pHluorin as a function of pH. Error bars represent SD of n=3 technical replicates.

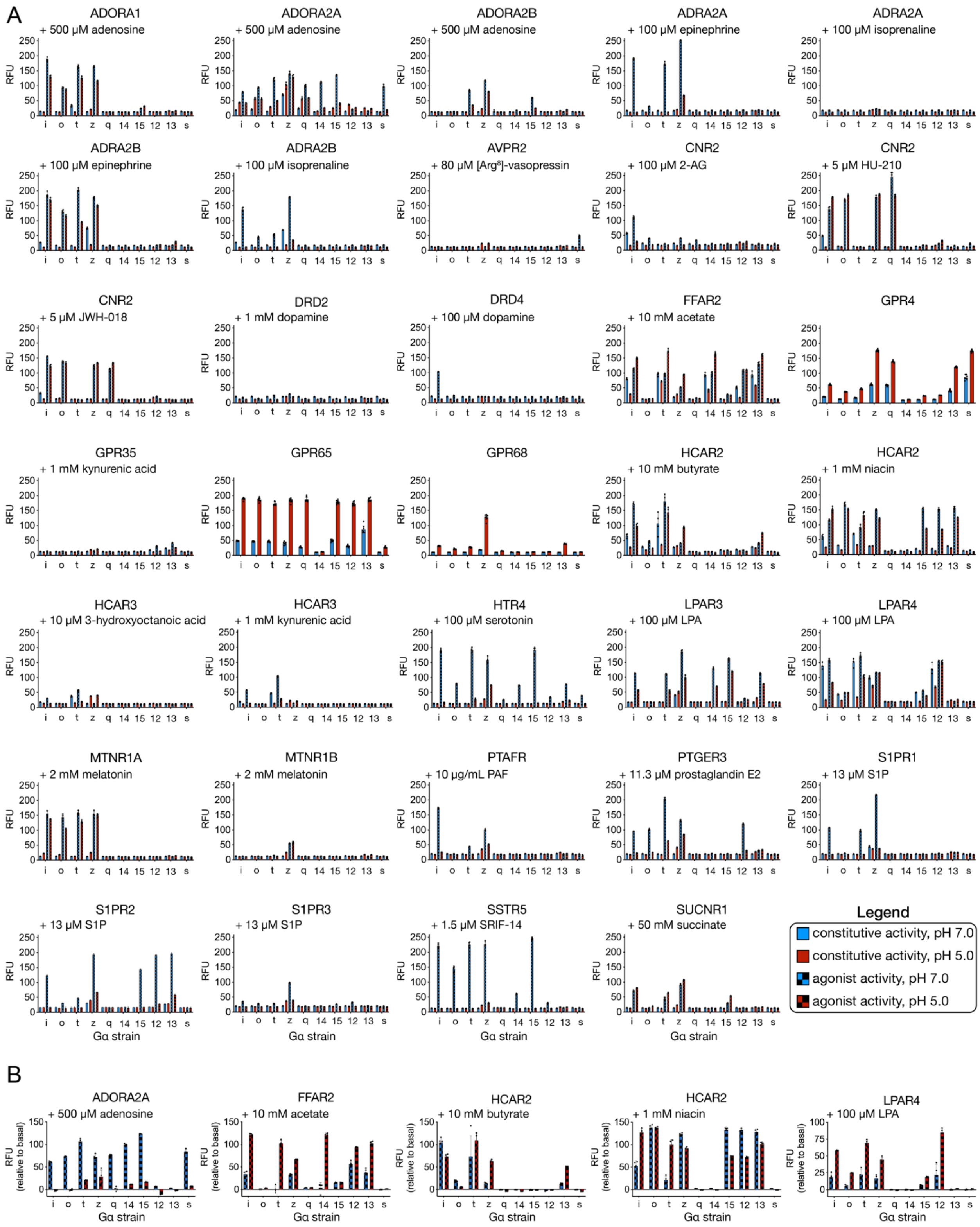
A
ADORA1
+ 500 µM adenosine
ADORA2A
+ 500 µM adenosine
ADORA2B
+ 500 µM adenosine
ADRA2A
+ 100 µM epinephrine
ADRA2A
+ 100 µM isoprenaline
ADRA2B
+ 100 µM epinephrine
ADRA2B
+ 100 µM isoprenaline
AVPR2
+ 80 µM [Arg8]-vasopressin
CNR2
+ 100 µM 2-AG
CNR2
+ 5 µM HU-210
CNR2
+ 5 µM JWH-018
DRD2
+ 1 mM dopamine
DRD4
+ 100 µM dopamine
FFAR2
+ 10 mM acetate
GPR4
GPR35
+ 1 mM kynurenic acid
GPR65
GPR68
HCAR2
+ 10 mM butyrate
HCAR2
+ 1 mM niacin
HCAR3
+ 10 µM 3-hydroxyoctanoic acid
HCAR3
+ 1 mM kynurenic acid
HTR4
+ 100 µM serotonin
LPAR3
+ 100 µM LPA
LPAR4
+ 100 µM LPA
MTNR1A
+ 2 mM melatonin
MTNR1B
+ 2 mM melatonin
PTAFR
+ 10 µg/mL PAF
PTGER3
+ 11.3 µM prostaglandin E2
S1PR1
+ 13 µM S1P
S1PR2
+ 13 µM S1P
S1PR3
+ 13 µM S1P
SSTR5
+ 1.5 µM SRIF-14
SUCNR1
+ 50 mM succinate
RFU
i o t z q 14 15 12 13 s
Gα strain
Legend
constitutive activity, pH 7.0
constitutive activity, pH 5.0
agonist activity, pH 7.0
agonist activity, pH 5.0
B
ADORA2A
+ 500 µM adenosine
FFAR2
+ 10 mM acetate
HCAR2
+ 10 mM butyrate
HCAR2
+ 1 mM niacin
LPAR4
+ 100 µM LPA
RFU
(relative to basal)
i o t z q 14 15 12 13 s
Gα strain

**Figure S2. DCyFIRscreen profiles of 280 human GPCR-Gα combinations as a function of pH (Related to Figs. 2-4).**

(A) Relative mTq2 fluorescence (RFU) of constitutive activity (solid bars) and agonist activity (checkered bars) for each GPCR in all 10 DCyFIR strains at pH 7 (blue) and 5 (red).

(B) RFU of treated relative to untreated cells for receptors with high levels of constitutive activity at pH 7 (blue checkered bars) and 5 (red checkered bars).

(A, B) Error bars represent SD of n=4 experimental replicates.

**Dataset S1. List of all yeast strains and GPCR names used in this study (Related to Figs. 1-4).**

| Strain name | Source | Abbreviated name | GPCR name | GPCR abbreviated name |
|---|---|---|---|---|
| BY4741 | Gift from Dohlman lab | BY4741 | N/A | N/A |
| BY4741 far1Δ sst2Δ fig1Δ::mTq2 | Kapolka et al., 2020 | DI2Δ fig1Δ::mTq2 | N/A | N/A |
| BY4741 far1Δ sst2Δ ste2Δ fig1Δ::mTq2 X-2:$P_{TEF1a}$-UnTS-$T_{CYC1b}$ gpa1(468-472)(KIGII>ECGLY) | Kapolka et al., 2020 | DI DCyFIR P1 I | N/A | N/A |
| BY4741 far1Δ sst2Δ ste2Δ fig1Δ::mTq2 X-2:$P_{TEF1a}$-UnTS-$T_{CYC1b}$ gpa1(468-472)(KIGII>GCGLY) | Kapolka et al., 2020 | DI DCyFIR P1 O | N/A | N/A |
| BY4741 far1Δ sst2Δ ste2Δ fig1Δ::mTq2 X-2:$P_{TEF1a}$-UnTS-$T_{CYC1b}$ gpa1(468-472)(KIGII>DCGLF) | Kapolka et al., 2020 | DI DCyFIR P1 T | N/A | N/A |
| BY4741 far1Δ sst2Δ ste2Δ fig1Δ::mTq2 X-2:$P_{TEF1a}$-UnTS-$T_{CYC1b}$ gpa1(468-472)(KIGII>YIGLC) | Kapolka et al., 2020 | DI DCyFIR P1 Z | N/A | N/A |
| BY4741 far1Δ sst2Δ ste2Δ fig1Δ::mTq2 X-2:$P_{TEF1a}$-UnTS-$T_{CYC1b}$ gpa1(468-472)(KIGII>EYNLV) | Kapolka et al., 2020 | DI DCyFIR P1 Q | N/A | N/A |
| BY4741 far1Δ sst2Δ ste2Δ fig1Δ::mTq2 X-2:$P_{TEF1a}$-UnTS-$T_{CYC1b}$ gpa1(468-472)(KIGII>EFNLV) | Kapolka et al., 2020 | DI DCyFIR P1 14 | N/A | N/A |
| BY4741 far1Δ sst2Δ ste2Δ fig1Δ::mTq2 X-2:$P_{TEF1a}$-UnTS-$T_{CYC1b}$ gpa1(468-472)(KIGII>EINLL) | Kapolka et al., 2020 | DI DCyFIR P1 15 | N/A | N/A |
| BY4741 far1Δ sst2Δ ste2Δ fig1Δ::mTq2 X-2:$P_{TEF1a}$-UnTS-$T_{CYC1b}$ gpa1(468-472)(KIGII>DIMLQ) | Kapolka et al., 2020 | DI DCyFIR P1 12 | N/A | N/A |
| BY4741 far1Δ sst2Δ ste2Δ fig1Δ::mTq2 X-2:$P_{TEF1a}$-UnTS-$T_{CYC1b}$ gpa1(468-472)(KIGII>QLMLQ) | Kapolka et al., 2020 | DI DCyFIR P1 13 | N/A | N/A |
| BY4741 far1Δ sst2Δ ste2Δ fig1Δ::mTq2 X-2:$P_{TEF1a}$-UnTS-$T_{CYC1b}$ gpa1(468-472)(KIGII>QYELL) | Kapolka et al., 2020 | DI DCyFIR P1 S | N/A | N/A |

| | | | | |
|---|---|---|---|---|
| BY4741 far1Δ sst2Δ ste2Δ fig1Δ::mTq2 X-2:$P_{TEF1a}$-ADORA1-$T_{CYC1b}$ gpa1(468-472)(KIGII>ECGLY) | Kapolka et al., 2020 | DI DCyFIR P1 I ADORA1 | adenosine A1 receptor | ADORA1 |
| BY4741 far1Δ sst2Δ ste2Δ fig1Δ::mTq2 X-2:$P_{TEF1a}$-ADORA1-$T_{CYC1b}$ gpa1(468-472)(KIGII>GCGLY) | Kapolka et al., 2020 | DI DCyFIR P1 O ADORA1 | adenosine A1 receptor | ADORA1 |
| BY4741 far1Δ sst2Δ ste2Δ fig1Δ::mTq2 X-2:$P_{TEF1a}$-ADORA1-$T_{CYC1b}$ gpa1(468-472)(KIGII>DCGLF) | Kapolka et al., 2020 | DI DCyFIR P1 T ADORA1 | adenosine A1 receptor | ADORA1 |
| BY4741 far1Δ sst2Δ ste2Δ fig1Δ::mTq2 X-2:$P_{TEF1a}$-ADORA1-$T_{CYC1b}$ gpa1(468-472)(KIGII>YIGLC) | Kapolka et al., 2020 | DI DCyFIR P1 Z ADORA1 | adenosine A1 receptor | ADORA1 |
| BY4741 far1Δ sst2Δ ste2Δ fig1Δ::mTq2 X-2:$P_{TEF1a}$-ADORA1-$T_{CYC1b}$ gpa1(468-472)(KIGII>EYNLV) | Kapolka et al., 2020 | DI DCyFIR P1 Q ADORA1 | adenosine A1 receptor | ADORA1 |
| BY4741 far1Δ sst2Δ ste2Δ fig1Δ::mTq2 X-2:$P_{TEF1a}$-ADORA1-$T_{CYC1b}$ gpa1(468-472)(KIGII>EFNLV) | Kapolka et al., 2020 | DI DCyFIR P1 14 ADORA1 | adenosine A1 receptor | ADORA1 |
| BY4741 far1Δ sst2Δ ste2Δ fig1Δ::mTq2 X-2:$P_{TEF1a}$-ADORA1-$T_{CYC1b}$ gpa1(468-472)(KIGII>EINLL) | Kapolka et al., 2020 | DI DCyFIR P1 15 ADORA1 | adenosine A1 receptor | ADORA1 |
| BY4741 far1Δ sst2Δ ste2Δ fig1Δ::mTq2 X-2:$P_{TEF1a}$-ADORA1-$T_{CYC1b}$ gpa1(468-472)(KIGII>DIMLQ) | Kapolka et al., 2020 | DI DCyFIR P1 12 ADORA1 | adenosine A1 receptor | ADORA1 |
| BY4741 far1Δ sst2Δ ste2Δ fig1Δ::mTq2 X-2:$P_{TEF1a}$-ADORA1-$T_{CYC1b}$ gpa1(468-472)(KIGII>QLMLQ) | Kapolka et al., 2020 | DI DCyFIR P1 13 ADORA1 | adenosine A1 receptor | ADORA1 |
| BY4741 far1Δ sst2Δ ste2Δ fig1Δ::mTq2 X-2:$P_{TEF1a}$-ADORA1-$T_{CYC1b}$ gpa1(468-472)(KIGII>QYELL) | Kapolka et al., 2020 | DI DCyFIR P1 S ADORA1 | adenosine A1 receptor | ADORA1 |
| BY4741 far1Δ sst2Δ ste2Δ fig1Δ::mTq2 X-2:$P_{TEF1a}$-ADORA2A-$T_{CYC1b}$ gpa1(468-472)(KIGII>ECGLY) | Kapolka et al., 2020 | DI DCyFIR P1 I ADORA2A | adenosine A2a receptor | ADORA2A |
| BY4741 far1Δ sst2Δ ste2Δ fig1Δ::mTq2 X-2:$P_{TEF1a}$-ADORA2A-$T_{CYC1b}$ gpa1(468-472)(KIGII>GCGLY) | Kapolka et al., 2020 | DI DCyFIR P1 O ADORA2A | adenosine A2a receptor | ADORA2A |
| BY4741 far1Δ sst2Δ ste2Δ fig1Δ::mTq2 X-2:$P_{TEF1a}$-ADORA2A-$T_{CYC1b}$ gpa1(468-472)(KIGII>DCGLF) | Kapolka et al., 2020 | DI DCyFIR P1 T ADORA2A | adenosine A2a receptor | ADORA2A |
| BY4741 far1Δ sst2Δ ste2Δ fig1Δ::mTq2 X-2:$P_{TEF1a}$-ADORA2A-$T_{CYC1b}$ gpa1(468-472)(KIGII>YIGLC) | Kapolka et al., 2020 | DI DCyFIR P1 Z ADORA2A | adenosine A2a receptor | ADORA2A |

| | | | | |
|---|---|---|---|---|
| BY4741 far1Δ sst2Δ ste2Δ fig1Δ::mTq2 X-2:$P_{TEF1a}$-ADORA2A-$T_{CYC1b}$ gpa1(468-472)(KIGII>EYNLV) | Kapolka et al., 2020 | DI DCyFIR P1 Q ADORA2A | adenosine A2a receptor | ADORA2A |
| BY4741 far1Δ sst2Δ ste2Δ fig1Δ::mTq2 X-2:$P_{TEF1a}$-ADORA2A-$T_{CYC1b}$ gpa1(468-472)(KIGII>EFNLV) | Kapolka et al., 2020 | DI DCyFIR P1 14 ADORA2A | adenosine A2a receptor | ADORA2A |
| BY4741 far1Δ sst2Δ ste2Δ fig1Δ::mTq2 X-2:$P_{TEF1a}$-ADORA2A-$T_{CYC1b}$ gpa1(468-472)(KIGII>EINLL) | Kapolka et al., 2020 | DI DCyFIR P1 15 ADORA2A | adenosine A2a receptor | ADORA2A |
| BY4741 far1Δ sst2Δ ste2Δ fig1Δ::mTq2 X-2:$P_{TEF1a}$-ADORA2A-$T_{CYC1b}$ gpa1(468-472)(KIGII>DIMLQ) | Kapolka et al., 2020 | DI DCyFIR P1 12 ADORA2A | adenosine A2a receptor | ADORA2A |
| BY4741 far1Δ sst2Δ ste2Δ fig1Δ::mTq2 X-2:$P_{TEF1a}$-ADORA2A-$T_{CYC1b}$ gpa1(468-472)(KIGII>QLMLQ) | Kapolka et al., 2020 | DI DCyFIR P1 13 ADORA2A | adenosine A2a receptor | ADORA2A |
| BY4741 far1Δ sst2Δ ste2Δ fig1Δ::mTq2 X-2:$P_{TEF1a}$-ADORA2A-$T_{CYC1b}$ gpa1(468-472)(KIGII>QYELL) | Kapolka et al., 2020 | DI DCyFIR P1 S ADORA2A | adenosine A2a receptor | ADORA2A |
| BY4741 far1Δ sst2Δ ste2Δ fig1Δ::mTq2 X-2:$P_{TEF1a}$-ADORA2B-$T_{CYC1b}$ gpa1(468-472)(KIGII>ECGLY) | Kapolka et al., 2020 | DI DCyFIR P1 I ADORA2B | adenosine A2b receptor | ADORA2B |
| BY4741 far1Δ sst2Δ ste2Δ fig1Δ::mTq2 X-2:$P_{TEF1a}$-ADORA2B-$T_{CYC1b}$ gpa1(468-472)(KIGII>GCGLY) | Kapolka et al., 2020 | DI DCyFIR P1 O ADORA2B | adenosine A2b receptor | ADORA2B |
| BY4741 far1Δ sst2Δ ste2Δ fig1Δ::mTq2 X-2:$P_{TEF1a}$-ADORA2B-$T_{CYC1b}$ gpa1(468-472)(KIGII>DCGLF) | Kapolka et al., 2020 | DI DCyFIR P1 T ADORA2B | adenosine A2b receptor | ADORA2B |
| BY4741 far1Δ sst2Δ ste2Δ fig1Δ::mTq2 X-2:$P_{TEF1a}$-ADORA2B-$T_{CYC1b}$ gpa1(468-472)(KIGII>YIGLC) | Kapolka et al., 2020 | DI DCyFIR P1 Z ADORA2B | adenosine A2b receptor | ADORA2B |
| BY4741 far1Δ sst2Δ ste2Δ fig1Δ::mTq2 X-2:$P_{TEF1a}$-ADORA2B-$T_{CYC1b}$ gpa1(468-472)(KIGII>EYNLV) | Kapolka et al., 2020 | DI DCyFIR P1 Q ADORA2B | adenosine A2b receptor | ADORA2B |
| BY4741 far1Δ sst2Δ ste2Δ fig1Δ::mTq2 X-2:$P_{TEF1a}$-ADORA2B-$T_{CYC1b}$ gpa1(468-472)(KIGII>EFNLV) | Kapolka et al., 2020 | DI DCyFIR P1 14 ADORA2B | adenosine A2b receptor | ADORA2B |
| BY4741 far1Δ sst2Δ ste2Δ fig1Δ::mTq2 X-2:$P_{TEF1a}$-ADORA2B-$T_{CYC1b}$ gpa1(468-472)(KIGII>EINLL) | Kapolka et al., 2020 | DI DCyFIR P1 15 ADORA2B | adenosine A2b receptor | ADORA2B |
| BY4741 far1Δ sst2Δ ste2Δ fig1Δ::mTq2 X-2:$P_{TEF1a}$-ADORA2B-$T_{CYC1b}$ gpa1(468-472)(KIGII>DIMLQ) | Kapolka et al., 2020 | DI DCyFIR P1 12 ADORA2B | adenosine A2b receptor | ADORA2B |

| | | | | |
|---|---|---|---|---|
| BY4741 far1Δ sst2Δ ste2Δ fig1Δ::mTq2 X-2:$P_{TEF1a}$-ADORA2B-$T_{CYC1b}$ gpa1(468-472)(KIGII>QLMLQ) | Kapolka et al., 2020 | DI DCyFIR P1 13 ADORA2B | adenosine A2b receptor | ADORA2B |
| BY4741 far1Δ sst2Δ ste2Δ fig1Δ::mTq2 X-2:$P_{TEF1a}$-ADORA2B-$T_{CYC1b}$ gpa1(468-472)(KIGII>QYELL) | Kapolka et al., 2020 | DI DCyFIR P1 S ADORA2B | adenosine A2b receptor | ADORA2B |
| BY4741 far1Δ sst2Δ ste2Δ fig1Δ::mTq2 X-2:$P_{TEF1a}$-ADRA2A-$T_{CYC1b}$ gpa1(468-472)(KIGII>ECGLY) | Kapolka et al., 2020 | DI DCyFIR P1 I ADRA2A | alpha-2A adrenergic receptor | ADRA2A |
| BY4741 far1Δ sst2Δ ste2Δ fig1Δ::mTq2 X-2:$P_{TEF1a}$-ADRA2A-$T_{CYC1b}$ gpa1(468-472)(KIGII>GCGLY) | Kapolka et al., 2020 | DI DCyFIR P1 O ADRA2A | alpha-2A adrenergic receptor | ADRA2A |
| BY4741 far1Δ sst2Δ ste2Δ fig1Δ::mTq2 X-2:$P_{TEF1a}$-ADRA2A-$T_{CYC1b}$ gpa1(468-472)(KIGII>DCGLF) | Kapolka et al., 2020 | DI DCyFIR P1 T ADRA2A | alpha-2A adrenergic receptor | ADRA2A |
| BY4741 far1Δ sst2Δ ste2Δ fig1Δ::mTq2 X-2:$P_{TEF1a}$-ADRA2A-$T_{CYC1b}$ gpa1(468-472)(KIGII>YIGLC) | Kapolka et al., 2020 | DI DCyFIR P1 Z ADRA2A | alpha-2A adrenergic receptor | ADRA2A |
| BY4741 far1Δ sst2Δ ste2Δ fig1Δ::mTq2 X-2:$P_{TEF1a}$-ADRA2A-$T_{CYC1b}$ gpa1(468-472)(KIGII>EYNLV) | Kapolka et al., 2020 | DI DCyFIR P1 Q ADRA2A | alpha-2A adrenergic receptor | ADRA2A |
| BY4741 far1Δ sst2Δ ste2Δ fig1Δ::mTq2 X-2:$P_{TEF1a}$-ADRA2A-$T_{CYC1b}$ gpa1(468-472)(KIGII>EFNLV) | Kapolka et al., 2020 | DI DCyFIR P1 14 ADRA2A | alpha-2A adrenergic receptor | ADRA2A |
| BY4741 far1Δ sst2Δ ste2Δ fig1Δ::mTq2 X-2:$P_{TEF1a}$-ADRA2A-$T_{CYC1b}$ gpa1(468-472)(KIGII>EINLL) | Kapolka et al., 2020 | DI DCyFIR P1 15 ADRA2A | alpha-2A adrenergic receptor | ADRA2A |
| BY4741 far1Δ sst2Δ ste2Δ fig1Δ::mTq2 X-2:$P_{TEF1a}$-ADRA2A-$T_{CYC1b}$ gpa1(468-472)(KIGII>DIMLQ) | Kapolka et al., 2020 | DI DCyFIR P1 12 ADRA2A | alpha-2A adrenergic receptor | ADRA2A |
| BY4741 far1Δ sst2Δ ste2Δ fig1Δ::mTq2 X-2:$P_{TEF1a}$-ADRA2A-$T_{CYC1b}$ gpa1(468-472)(KIGII>QLMLQ) | Kapolka et al., 2020 | DI DCyFIR P1 13 ADRA2A | alpha-2A adrenergic receptor | ADRA2A |
| BY4741 far1Δ sst2Δ ste2Δ fig1Δ::mTq2 X-2:$P_{TEF1a}$-ADRA2A-$T_{CYC1b}$ gpa1(468-472)(KIGII>QYELL) | Kapolka et al., 2020 | DI DCyFIR P1 S ADRA2A | alpha-2A adrenergic receptor | ADRA2A |
| BY4741 far1Δ sst2Δ ste2Δ fig1Δ::mTq2 X-2:$P_{TEF1a}$-ADRA2B-$T_{CYC1b}$ gpa1(468-472)(KIGII>ECGLY) | Kapolka et al., 2020 | DI DCyFIR P1 I ADRA2B | alpha-2B adrenergic receptor | ADRA2B |
| BY4741 far1Δ sst2Δ ste2Δ fig1Δ::mTq2 X-2:$P_{TEF1a}$-ADRA2B-$T_{CYC1b}$ gpa1(468-472)(KIGII>GCGLY) | Kapolka et al., 2020 | DI DCyFIR P1 O ADRA2B | alpha-2B adrenergic receptor | ADRA2B |

| | | | | |
|---|---|---|---|---|
| BY4741 far1Δ sst2Δ ste2Δ fig1Δ::mTq2 X-2:$P_{TEF1a}$-ADRA2B-$T_{CYC1b}$ gpa1(468-472)(KIGII>DCGLF) | Kapolka et al., 2020 | DI DCyFIR P1 T ADRA2B | alpha-2B adrenergic receptor | ADRA2B |
| BY4741 far1Δ sst2Δ ste2Δ fig1Δ::mTq2 X-2:$P_{TEF1a}$-ADRA2B-$T_{CYC1b}$ gpa1(468-472)(KIGII>YIGLC) | Kapolka et al., 2020 | DI DCyFIR P1 Z ADRA2B | alpha-2B adrenergic receptor | ADRA2B |
| BY4741 far1Δ sst2Δ ste2Δ fig1Δ::mTq2 X-2:$P_{TEF1a}$-ADRA2B-$T_{CYC1b}$ gpa1(468-472)(KIGII>EYNLV) | Kapolka et al., 2020 | DI DCyFIR P1 Q ADRA2B | alpha-2B adrenergic receptor | ADRA2B |
| BY4741 far1Δ sst2Δ ste2Δ fig1Δ::mTq2 X-2:$P_{TEF1a}$-ADRA2B-$T_{CYC1b}$ gpa1(468-472)(KIGII>EFNLV) | Kapolka et al., 2020 | DI DCyFIR P1 14 ADRA2B | alpha-2B adrenergic receptor | ADRA2B |
| BY4741 far1Δ sst2Δ ste2Δ fig1Δ::mTq2 X-2:$P_{TEF1a}$-ADRA2B-$T_{CYC1b}$ gpa1(468-472)(KIGII>EINLL) | Kapolka et al., 2020 | DI DCyFIR P1 15 ADRA2B | alpha-2B adrenergic receptor | ADRA2B |
| BY4741 far1Δ sst2Δ ste2Δ fig1Δ::mTq2 X-2:$P_{TEF1a}$-ADRA2B-$T_{CYC1b}$ gpa1(468-472)(KIGII>DIMLQ) | Kapolka et al., 2020 | DI DCyFIR P1 12 ADRA2B | alpha-2B adrenergic receptor | ADRA2B |
| BY4741 far1Δ sst2Δ ste2Δ fig1Δ::mTq2 X-2:$P_{TEF1a}$-ADRA2B-$T_{CYC1b}$ gpa1(468-472)(KIGII>QLMLQ) | Kapolka et al., 2020 | DI DCyFIR P1 13 ADRA2B | alpha-2B adrenergic receptor | ADRA2B |
| BY4741 far1Δ sst2Δ ste2Δ fig1Δ::mTq2 X-2:$P_{TEF1a}$-ADRA2B-$T_{CYC1b}$ gpa1(468-472)(KIGII>QYELL) | Kapolka et al., 2020 | DI DCyFIR P1 S ADRA2B | alpha-2B adrenergic receptor | ADRA2B |
| BY4741 far1Δ sst2Δ ste2Δ fig1Δ::mTq2 X-2:$P_{TEF1a}$-AVPR2-$T_{CYC1b}$ gpa1(468-472)(KIGII>ECGLY) | Kapolka et al., 2020 | DI DCyFIR P1 I AVPR2 | arginine vasopressin receptor 2 | AVPR2 |
| BY4741 far1Δ sst2Δ ste2Δ fig1Δ::mTq2 X-2:$P_{TEF1a}$-AVPR2-$T_{CYC1b}$ gpa1(468-472)(KIGII>GCGLY) | Kapolka et al., 2020 | DI DCyFIR P1 O AVPR2 | arginine vasopressin receptor 2 | AVPR2 |
| BY4741 far1Δ sst2Δ ste2Δ fig1Δ::mTq2 X-2:$P_{TEF1a}$-AVPR2-$T_{CYC1b}$ gpa1(468-472)(KIGII>DCGLF) | Kapolka et al., 2020 | DI DCyFIR P1 T AVPR2 | arginine vasopressin receptor 2 | AVPR2 |
| BY4741 far1Δ sst2Δ ste2Δ fig1Δ::mTq2 X-2:$P_{TEF1a}$-AVPR2-$T_{CYC1b}$ gpa1(468-472)(KIGII>YIGLC) | Kapolka et al., 2020 | DI DCyFIR P1 Z AVPR2 | arginine vasopressin receptor 2 | AVPR2 |
| BY4741 far1Δ sst2Δ ste2Δ fig1Δ::mTq2 X-2:$P_{TEF1a}$-AVPR2-$T_{CYC1b}$ gpa1(468-472)(KIGII>EYNLV) | Kapolka et al., 2020 | DI DCyFIR P1 Q AVPR2 | arginine vasopressin receptor 2 | AVPR2 |
| BY4741 far1Δ sst2Δ ste2Δ fig1Δ::mTq2 X-2:$P_{TEF1a}$-AVPR2-$T_{CYC1b}$ gpa1(468-472)(KIGII>EFNLV) | Kapolka et al., 2020 | DI DCyFIR P1 14 AVPR2 | arginine vasopressin receptor 2 | AVPR2 |

| | | | | |
|---|---|---|---|---|
| BY4741 far1Δ sst2Δ ste2Δ fig1Δ::mTq2 X-2:$P_{TEF1a}$-AVPR2-$T_{CYC1b}$ gpa1(468-472)(KIGII>EINLL) | Kapolka et al., 2020 | DI DCyFIR P1 15 AVPR2 | arginine vasopressin receptor 2 | AVPR2 |
| BY4741 far1Δ sst2Δ ste2Δ fig1Δ::mTq2 X-2:$P_{TEF1a}$-AVPR2-$T_{CYC1b}$ gpa1(468-472)(KIGII>DIMLQ) | Kapolka et al., 2020 | DI DCyFIR P1 12 AVPR2 | arginine vasopressin receptor 2 | AVPR2 |
| BY4741 far1Δ sst2Δ ste2Δ fig1Δ::mTq2 X-2:$P_{TEF1a}$-AVPR2-$T_{CYC1b}$ gpa1(468-472)(KIGII>QLMLQ) | Kapolka et al., 2020 | DI DCyFIR P1 13 AVPR2 | arginine vasopressin receptor 2 | AVPR2 |
| BY4741 far1Δ sst2Δ ste2Δ fig1Δ::mTq2 X-2:$P_{TEF1a}$-AVPR2-$T_{CYC1b}$ gpa1(468-472)(KIGII>QYELL) | Kapolka et al., 2020 | DI DCyFIR P1 S AVPR2 | arginine vasopressin receptor 2 | AVPR2 |
| BY4741 far1Δ sst2Δ ste2Δ fig1Δ::mTq2 X-2:$P_{TEF1a}$-CNR2-$T_{CYC1b}$ gpa1(468-472)(KIGII>ECGLY) | Kapolka et al., 2020 | DI DCyFIR P1 I CNR2 | cannabinoid receptor 2 | CNR2 |
| BY4741 far1Δ sst2Δ ste2Δ fig1Δ::mTq2 X-2:$P_{TEF1a}$-CNR2-$T_{CYC1b}$ gpa1(468-472)(KIGII>GCGLY) | Kapolka et al., 2020 | DI DCyFIR P1 O CNR2 | cannabinoid receptor 2 | CNR2 |
| BY4741 far1Δ sst2Δ ste2Δ fig1Δ::mTq2 X-2:$P_{TEF1a}$-CNR2-$T_{CYC1b}$ gpa1(468-472)(KIGII>DCGLF) | Kapolka et al., 2020 | DI DCyFIR P1 T CNR2 | cannabinoid receptor 2 | CNR2 |
| BY4741 far1Δ sst2Δ ste2Δ fig1Δ::mTq2 X-2:$P_{TEF1a}$-CNR2-$T_{CYC1b}$ gpa1(468-472)(KIGII>YIGLC) | Kapolka et al., 2020 | DI DCyFIR P1 Z CNR2 | cannabinoid receptor 2 | CNR2 |
| BY4741 far1Δ sst2Δ ste2Δ fig1Δ::mTq2 X-2:$P_{TEF1a}$-CNR2-$T_{CYC1b}$ gpa1(468-472)(KIGII>EYNLV) | Kapolka et al., 2020 | DI DCyFIR P1 Q CNR2 | cannabinoid receptor 2 | CNR2 |
| BY4741 far1Δ sst2Δ ste2Δ fig1Δ::mTq2 X-2:$P_{TEF1a}$-CNR2-$T_{CYC1b}$ gpa1(468-472)(KIGII>EFNLV) | Kapolka et al., 2020 | DI DCyFIR P1 14 CNR2 | cannabinoid receptor 2 | CNR2 |
| BY4741 far1Δ sst2Δ ste2Δ fig1Δ::mTq2 X-2:$P_{TEF1a}$-CNR2-$T_{CYC1b}$ gpa1(468-472)(KIGII>EINLL) | Kapolka et al., 2020 | DI DCyFIR P1 15 CNR2 | cannabinoid receptor 2 | CNR2 |
| BY4741 far1Δ sst2Δ ste2Δ fig1Δ::mTq2 X-2:$P_{TEF1a}$-CNR2-$T_{CYC1b}$ gpa1(468-472)(KIGII>DIMLQ) | Kapolka et al., 2020 | DI DCyFIR P1 12 CNR2 | cannabinoid receptor 2 | CNR2 |
| BY4741 far1Δ sst2Δ ste2Δ fig1Δ::mTq2 X-2:$P_{TEF1a}$-CNR2-$T_{CYC1b}$ gpa1(468-472)(KIGII>QLMLQ) | Kapolka et al., 2020 | DI DCyFIR P1 13 CNR2 | cannabinoid receptor 2 | CNR2 |
| BY4741 far1Δ sst2Δ ste2Δ fig1Δ::mTq2 X-2:$P_{TEF1a}$-CNR2-$T_{CYC1b}$ gpa1(468-472)(KIGII>QYELL) | Kapolka et al., 2020 | DI DCyFIR P1 S CNR2 | cannabinoid receptor 2 | CNR2 |

| | | | | |
|---|---|---|---|---|
| BY4741 far1Δ sst2Δ ste2Δ fig1Δ::mTq2 X-2:$P_{TEF1a}$-DRD2-$T_{CYC1b}$ gpa1(468-472)(KIGII>ECGLY) | This paper | DI DCyFIR P1 I DRD2 | dopamine receptor D2 | DRD2 |
| BY4741 far1Δ sst2Δ ste2Δ fig1Δ::mTq2 X-2:$P_{TEF1a}$-DRD2-$T_{CYC1b}$ gpa1(468-472)(KIGII>GCGLY) | This paper | DI DCyFIR P1 O DRD2 | dopamine receptor D2 | DRD2 |
| BY4741 far1Δ sst2Δ ste2Δ fig1Δ::mTq2 X-2:$P_{TEF1a}$-DRD2-$T_{CYC1b}$ gpa1(468-472)(KIGII>DCGLF) | This paper | DI DCyFIR P1 T DRD2 | dopamine receptor D2 | DRD2 |
| BY4741 far1Δ sst2Δ ste2Δ fig1Δ::mTq2 X-2:$P_{TEF1a}$-DRD2-$T_{CYC1b}$ gpa1(468-472)(KIGII>YIGLC) | This paper | DI DCyFIR P1 Z DRD2 | dopamine receptor D2 | DRD2 |
| BY4741 far1Δ sst2Δ ste2Δ fig1Δ::mTq2 X-2:$P_{TEF1a}$-DRD2-$T_{CYC1b}$ gpa1(468-472)(KIGII>EYNLV) | This paper | DI DCyFIR P1 Q DRD2 | dopamine receptor D2 | DRD2 |
| BY4741 far1Δ sst2Δ ste2Δ fig1Δ::mTq2 X-2:$P_{TEF1a}$-DRD2-$T_{CYC1b}$ gpa1(468-472)(KIGII>EFNLV) | This paper | DI DCyFIR P1 14 DRD2 | dopamine receptor D2 | DRD2 |
| BY4741 far1Δ sst2Δ ste2Δ fig1Δ::mTq2 X-2:$P_{TEF1a}$-DRD2-$T_{CYC1b}$ gpa1(468-472)(KIGII>EINLL) | This paper | DI DCyFIR P1 15 DRD2 | dopamine receptor D2 | DRD2 |
| BY4741 far1Δ sst2Δ ste2Δ fig1Δ::mTq2 X-2:$P_{TEF1a}$-DRD2-$T_{CYC1b}$ gpa1(468-472)(KIGII>DIMLQ) | This paper | DI DCyFIR P1 12 DRD2 | dopamine receptor D2 | DRD2 |
| BY4741 far1Δ sst2Δ ste2Δ fig1Δ::mTq2 X-2:$P_{TEF1a}$-DRD2-$T_{CYC1b}$ gpa1(468-472)(KIGII>QLMLQ) | This paper | DI DCyFIR P1 13 DRD2 | dopamine receptor D2 | DRD2 |
| BY4741 far1Δ sst2Δ ste2Δ fig1Δ::mTq2 X-2:$P_{TEF1a}$-DRD2-$T_{CYC1b}$ gpa1(468-472)(KIGII>QYELL) | This paper | DI DCyFIR P1 S DRD2 | dopamine receptor D2 | DRD2 |
| BY4741 far1Δ sst2Δ ste2Δ fig1Δ::mTq2 X-2:$P_{TEF1a}$-DRD4-$T_{CYC1b}$ gpa1(468-472)(KIGII>ECGLY) | This paper | DI DCyFIR P1 I DRD4 | dopamine receptor D4 | DRD4 |
| BY4741 far1Δ sst2Δ ste2Δ fig1Δ::mTq2 X-2:$P_{TEF1a}$-DRD4-$T_{CYC1b}$ gpa1(468-472)(KIGII>GCGLY) | This paper | DI DCyFIR P1 O DRD4 | dopamine receptor D4 | DRD4 |
| BY4741 far1Δ sst2Δ ste2Δ fig1Δ::mTq2 X-2:$P_{TEF1a}$-DRD4-$T_{CYC1b}$ gpa1(468-472)(KIGII>DCGLF) | This paper | DI DCyFIR P1 T DRD4 | dopamine receptor D4 | DRD4 |
| BY4741 far1Δ sst2Δ ste2Δ fig1Δ::mTq2 X-2:$P_{TEF1a}$-DRD4-$T_{CYC1b}$ gpa1(468-472)(KIGII>YIGLC) | This paper | DI DCyFIR P1 Z DRD4 | dopamine receptor D4 | DRD4 |

| | | | | |
|---|---|---|---|---|
| BY4741 far1Δ sst2Δ ste2Δ fig1Δ::mTq2 X-2:P$_{TEF1a}$-DRD4-T$_{CYC1b}$ gpa1(468-472)(KIGII>EYNLV) | This paper | DI DCyFIR P1 Q DRD4 | dopamine receptor D4 | DRD4 |
| BY4741 far1Δ sst2Δ ste2Δ fig1Δ::mTq2 X-2:P$_{TEF1a}$-DRD4-T$_{CYC1b}$ gpa1(468-472)(KIGII>EFNLV) | This paper | DI DCyFIR P1 14 DRD4 | dopamine receptor D4 | DRD4 |
| BY4741 far1Δ sst2Δ ste2Δ fig1Δ::mTq2 X-2:P$_{TEF1a}$-DRD4-T$_{CYC1b}$ gpa1(468-472)(KIGII>EINLL) | This paper | DI DCyFIR P1 15 DRD4 | dopamine receptor D4 | DRD4 |
| BY4741 far1Δ sst2Δ ste2Δ fig1Δ::mTq2 X-2:P$_{TEF1a}$-DRD4-T$_{CYC1b}$ gpa1(468-472)(KIGII>DIMLQ) | This paper | DI DCyFIR P1 12 DRD4 | dopamine receptor D4 | DRD4 |
| BY4741 far1Δ sst2Δ ste2Δ fig1Δ::mTq2 X-2:P$_{TEF1a}$-DRD4-T$_{CYC1b}$ gpa1(468-472)(KIGII>QLMLQ) | This paper | DI DCyFIR P1 13 DRD4 | dopamine receptor D4 | DRD4 |
| BY4741 far1Δ sst2Δ ste2Δ fig1Δ::mTq2 X-2:P$_{TEF1a}$-DRD4-T$_{CYC1b}$ gpa1(468-472)(KIGII>QYELL) | This paper | DI DCyFIR P1 S DRD4 | dopamine receptor D4 | DRD4 |
| BY4741 far1Δ sst2Δ ste2Δ fig1Δ::mTq2 X-2:P$_{TEF1a}$-FFAR2-T$_{CYC1b}$ gpa1(468-472)(KIGII>ECGLY) | Kapolka et al., 2020 | DI DCyFIR P1 I FFAR2 | free fatty acid receptor 2 | FFAR2 |
| BY4741 far1Δ sst2Δ ste2Δ fig1Δ::mTq2 X-2:P$_{TEF1a}$-FFAR2-T$_{CYC1b}$ gpa1(468-472)(KIGII>GCGLY) | Kapolka et al., 2020 | DI DCyFIR P1 O FFAR2 | free fatty acid receptor 2 | FFAR2 |
| BY4741 far1Δ sst2Δ ste2Δ fig1Δ::mTq2 X-2:P$_{TEF1a}$-FFAR2-T$_{CYC1b}$ gpa1(468-472)(KIGII>DCGLF) | Kapolka et al., 2020 | DI DCyFIR P1 T FFAR2 | free fatty acid receptor 2 | FFAR2 |
| BY4741 far1Δ sst2Δ ste2Δ fig1Δ::mTq2 X-2:P$_{TEF1a}$-FFAR2-T$_{CYC1b}$ gpa1(468-472)(KIGII>YIGLC) | Kapolka et al., 2020 | DI DCyFIR P1 Z FFAR2 | free fatty acid receptor 2 | FFAR2 |
| BY4741 far1Δ sst2Δ ste2Δ fig1Δ::mTq2 X-2:P$_{TEF1a}$-FFAR2-T$_{CYC1b}$ gpa1(468-472)(KIGII>EYNLV) | Kapolka et al., 2020 | DI DCyFIR P1 Q FFAR2 | free fatty acid receptor 2 | FFAR2 |
| BY4741 far1Δ sst2Δ ste2Δ fig1Δ::mTq2 X-2:P$_{TEF1a}$-FFAR2-T$_{CYC1b}$ gpa1(468-472)(KIGII>EFNLV) | Kapolka et al., 2020 | DI DCyFIR P1 14 FFAR2 | free fatty acid receptor 2 | FFAR2 |
| BY4741 far1Δ sst2Δ ste2Δ fig1Δ::mTq2 X-2:P$_{TEF1a}$-FFAR2-T$_{CYC1b}$ gpa1(468-472)(KIGII>EINLL) | Kapolka et al., 2020 | DI DCyFIR P1 15 FFAR2 | free fatty acid receptor 2 | FFAR2 |
| BY4741 far1Δ sst2Δ ste2Δ fig1Δ::mTq2 X-2:P$_{TEF1a}$-FFAR2-T$_{CYC1b}$ gpa1(468-472)(KIGII>DIMLQ) | Kapolka et al., 2020 | DI DCyFIR P1 12 FFAR2 | free fatty acid receptor 2 | FFAR2 |

| | | | | |
|---|---|---|---|---|
| BY4741 far1Δ sst2Δ ste2Δ fig1Δ::mTq2 X-2:$P_{TEF1a}$-FFAR2-$T_{CYC1b}$ gpa1(468-472)(KIGII>QLMLQ) | Kapolka et al., 2020 | DI DCyFIR P1 13 FFAR2 | free fatty acid receptor 2 | FFAR2 |
| BY4741 far1Δ sst2Δ ste2Δ fig1Δ::mTq2 X-2:$P_{TEF1a}$-FFAR2-$T_{CYC1b}$ gpa1(468-472)(KIGII>QYELL) | Kapolka et al., 2020 | DI DCyFIR P1 S FFAR2 | free fatty acid receptor 2 | FFAR2 |
| BY4741 far1Δ sst2Δ ste2Δ fig1Δ::mTq2 X-2:$P_{TEF1a}$-GPR35-$T_{CYC1b}$ gpa1(468-472)(KIGII>ECGLY) | Kapolka et al., 2020 | DI DCyFIR P1 I GPR35 | G protein-coupled receptor 35 | GPR35 |
| BY4741 far1Δ sst2Δ ste2Δ fig1Δ::mTq2 X-2:$P_{TEF1a}$-GPR35-$T_{CYC1b}$ gpa1(468-472)(KIGII>GCGLY) | Kapolka et al., 2020 | DI DCyFIR P1 O GPR35 | G protein-coupled receptor 35 | GPR35 |
| BY4741 far1Δ sst2Δ ste2Δ fig1Δ::mTq2 X-2:$P_{TEF1a}$-GPR35-$T_{CYC1b}$ gpa1(468-472)(KIGII>DCGLF) | Kapolka et al., 2020 | DI DCyFIR P1 T GPR35 | G protein-coupled receptor 35 | GPR35 |
| BY4741 far1Δ sst2Δ ste2Δ fig1Δ::mTq2 X-2:$P_{TEF1a}$-GPR35-$T_{CYC1b}$ gpa1(468-472)(KIGII>YIGLC) | Kapolka et al., 2020 | DI DCyFIR P1 Z GPR35 | G protein-coupled receptor 35 | GPR35 |
| BY4741 far1Δ sst2Δ ste2Δ fig1Δ::mTq2 X-2:$P_{TEF1a}$-GPR35-$T_{CYC1b}$ gpa1(468-472)(KIGII>EYNLV) | Kapolka et al., 2020 | DI DCyFIR P1 Q GPR35 | G protein-coupled receptor 35 | GPR35 |
| BY4741 far1Δ sst2Δ ste2Δ fig1Δ::mTq2 X-2:$P_{TEF1a}$-GPR35-$T_{CYC1b}$ gpa1(468-472)(KIGII>EFNLV) | Kapolka et al., 2020 | DI DCyFIR P1 14 GPR35 | G protein-coupled receptor 35 | GPR35 |
| BY4741 far1Δ sst2Δ ste2Δ fig1Δ::mTq2 X-2:$P_{TEF1a}$-GPR35-$T_{CYC1b}$ gpa1(468-472)(KIGII>EINLL) | Kapolka et al., 2020 | DI DCyFIR P1 15 GPR35 | G protein-coupled receptor 35 | GPR35 |
| BY4741 far1Δ sst2Δ ste2Δ fig1Δ::mTq2 X-2:$P_{TEF1a}$-GPR35-$T_{CYC1b}$ gpa1(468-472)(KIGII>DIMLQ) | Kapolka et al., 2020 | DI DCyFIR P1 12 GPR35 | G protein-coupled receptor 35 | GPR35 |
| BY4741 far1Δ sst2Δ ste2Δ fig1Δ::mTq2 X-2:$P_{TEF1a}$-GPR35-$T_{CYC1b}$ gpa1(468-472)(KIGII>QLMLQ) | Kapolka et al., 2020 | DI DCyFIR P1 13 GPR35 | G protein-coupled receptor 35 | GPR35 |
| BY4741 far1Δ sst2Δ ste2Δ fig1Δ::mTq2 X-2:$P_{TEF1a}$-GPR35-$T_{CYC1b}$ gpa1(468-472)(KIGII>QYELL) | Kapolka et al., 2020 | DI DCyFIR P1 S GPR35 | G protein-coupled receptor 35 | GPR35 |
| BY4741 far1Δ sst2Δ ste2Δ fig1Δ::mTq2 X-2:$P_{TEF1a}$-GPR4-$T_{CYC1b}$ gpa1(468-472)(KIGII>ECGLY) | Kapolka et al., 2020 | DI DCyFIR P1 I GPR4 | G protein-coupled receptor 4 | GPR4 |
| BY4741 far1Δ sst2Δ ste2Δ fig1Δ::mTq2 X-2:$P_{TEF1a}$-GPR4-$T_{CYC1b}$ gpa1(468-472)(KIGII>GCGLY) | Kapolka et al., 2020 | DI DCyFIR P1 O GPR4 | G protein-coupled receptor 4 | GPR4 |

| | | | | |
|---|---|---|---|---|
| BY4741 far1Δ sst2Δ ste2Δ fig1Δ::mTq2 X-2:$P_{TEF1a}$-GPR4-$T_{CYC1b}$ gpa1(468-472)(KIGII>DCGLF) | Kapolka et al., 2020 | DI DCyFIR P1 T GPR4 | G protein-coupled receptor 4 | GPR4 |
| BY4741 far1Δ sst2Δ ste2Δ fig1Δ::mTq2 X-2:$P_{TEF1a}$-GPR4-$T_{CYC1b}$ gpa1(468-472)(KIGII>YIGLC) | Kapolka et al., 2020 | DI DCyFIR P1 Z GPR4 | G protein-coupled receptor 4 | GPR4 |
| BY4741 far1Δ sst2Δ ste2Δ fig1Δ::mTq2 X-2:$P_{TEF1a}$-GPR4-$T_{CYC1b}$ gpa1(468-472)(KIGII>EYNLV) | Kapolka et al., 2020 | DI DCyFIR P1 Q GPR4 | G protein-coupled receptor 4 | GPR4 |
| BY4741 far1Δ sst2Δ ste2Δ fig1Δ::mTq2 X-2:$P_{TEF1a}$-GPR4-$T_{CYC1b}$ gpa1(468-472)(KIGII>EFNLV) | Kapolka et al., 2020 | DI DCyFIR P1 14 GPR4 | G protein-coupled receptor 4 | GPR4 |
| BY4741 far1Δ sst2Δ ste2Δ fig1Δ::mTq2 X-2:$P_{TEF1a}$-GPR4-$T_{CYC1b}$ gpa1(468-472)(KIGII>EINLL) | Kapolka et al., 2020 | DI DCyFIR P1 15 GPR4 | G protein-coupled receptor 4 | GPR4 |
| BY4741 far1Δ sst2Δ ste2Δ fig1Δ::mTq2 X-2:$P_{TEF1a}$-GPR4-$T_{CYC1b}$ gpa1(468-472)(KIGII>DIMLQ) | Kapolka et al., 2020 | DI DCyFIR P1 12 GPR4 | G protein-coupled receptor 4 | GPR4 |
| BY4741 far1Δ sst2Δ ste2Δ fig1Δ::mTq2 X-2:$P_{TEF1a}$-GPR4-$T_{CYC1b}$ gpa1(468-472)(KIGII>QLMLQ) | Kapolka et al., 2020 | DI DCyFIR P1 13 GPR4 | G protein-coupled receptor 4 | GPR4 |
| BY4741 far1Δ sst2Δ ste2Δ fig1Δ::mTq2 X-2:$P_{TEF1a}$-GPR4-$T_{CYC1b}$ gpa1(468-472)(KIGII>QYELL) | Kapolka et al., 2020 | DI DCyFIR P1 S GPR4 | G protein-coupled receptor 4 | GPR4 |
| BY4741 far1Δ sst2Δ ste2Δ fig1Δ::mTq2 X-2:$P_{TEF1a}$-GPR65-$T_{CYC1b}$ gpa1(468-472)(KIGII>ECGLY) | Kapolka et al., 2020 | DI DCyFIR P1 I GPR65 | G protein-coupled receptor 65 | GPR65 |
| BY4741 far1Δ sst2Δ ste2Δ fig1Δ::mTq2 X-2:$P_{TEF1a}$-GPR65-$T_{CYC1b}$ gpa1(468-472)(KIGII>GCGLY) | Kapolka et al., 2020 | DI DCyFIR P1 O GPR65 | G protein-coupled receptor 65 | GPR65 |
| BY4741 far1Δ sst2Δ ste2Δ fig1Δ::mTq2 X-2:$P_{TEF1a}$-GPR65-$T_{CYC1b}$ gpa1(468-472)(KIGII>DCGLF) | Kapolka et al., 2020 | DI DCyFIR P1 T GPR65 | G protein-coupled receptor 65 | GPR65 |
| BY4741 far1Δ sst2Δ ste2Δ fig1Δ::mTq2 X-2:$P_{TEF1a}$-GPR65-$T_{CYC1b}$ gpa1(468-472)(KIGII>YIGLC) | Kapolka et al., 2020 | DI DCyFIR P1 Z GPR65 | G protein-coupled receptor 65 | GPR65 |
| BY4741 far1Δ sst2Δ ste2Δ fig1Δ::mTq2 X-2:$P_{TEF1a}$-GPR65-$T_{CYC1b}$ gpa1(468-472)(KIGII>EYNLV) | Kapolka et al., 2020 | DI DCyFIR P1 Q GPR65 | G protein-coupled receptor 65 | GPR65 |
| BY4741 far1Δ sst2Δ ste2Δ fig1Δ::mTq2 X-2:$P_{TEF1a}$-GPR65-$T_{CYC1b}$ gpa1(468-472)(KIGII>EFNLV) | Kapolka et al., 2020 | DI DCyFIR P1 14 GPR65 | G protein-coupled receptor 65 | GPR65 |

| | | | | |
|---|---|---|---|---|
| BY4741 far1Δ sst2Δ ste2Δ fig1Δ::mTq2 X-2:$P_{TEF1a}$-GPR65-$T_{CYC1b}$ gpa1(468-472)(KIGII>EINLL) | Kapolka et al., 2020 | DI DCyFIR P1 15 GPR65 | G protein-coupled receptor 65 | GPR65 |
| BY4741 far1Δ sst2Δ ste2Δ fig1Δ::mTq2 X-2:$P_{TEF1a}$-GPR65-$T_{CYC1b}$ gpa1(468-472)(KIGII>DIMLQ) | Kapolka et al., 2020 | DI DCyFIR P1 12 GPR65 | G protein-coupled receptor 65 | GPR65 |
| BY4741 far1Δ sst2Δ ste2Δ fig1Δ::mTq2 X-2:$P_{TEF1a}$-GPR65-$T_{CYC1b}$ gpa1(468-472)(KIGII>QLMLQ) | Kapolka et al., 2020 | DI DCyFIR P1 13 GPR65 | G protein-coupled receptor 65 | GPR65 |
| BY4741 far1Δ sst2Δ ste2Δ fig1Δ::mTq2 X-2:$P_{TEF1a}$-GPR65-$T_{CYC1b}$ gpa1(468-472)(KIGII>QYELL) | Kapolka et al., 2020 | DI DCyFIR P1 S GPR65 | G protein-coupled receptor 65 | GPR65 |
| BY4741 far1Δ sst2Δ ste2Δ fig1Δ::mTq2 X-2:$P_{TEF1a}$-GPR68-$T_{CYC1b}$ gpa1(468-472)(KIGII>ECGLY) | Kapolka et al., 2020 | DI DCyFIR P1 I GPR68 | G protein-coupled receptor 68 | GPR68 |
| BY4741 far1Δ sst2Δ ste2Δ fig1Δ::mTq2 X-2:$P_{TEF1a}$-GPR68-$T_{CYC1b}$ gpa1(468-472)(KIGII>GCGLY) | Kapolka et al., 2020 | DI DCyFIR P1 O GPR68 | G protein-coupled receptor 68 | GPR68 |
| BY4741 far1Δ sst2Δ ste2Δ fig1Δ::mTq2 X-2:$P_{TEF1a}$-GPR68-$T_{CYC1b}$ gpa1(468-472)(KIGII>DCGLF) | Kapolka et al., 2020 | DI DCyFIR P1 T GPR68 | G protein-coupled receptor 68 | GPR68 |
| BY4741 far1Δ sst2Δ ste2Δ fig1Δ::mTq2 X-2:$P_{TEF1a}$-GPR68-$T_{CYC1b}$ gpa1(468-472)(KIGII>YIGLC) | Kapolka et al., 2020 | DI DCyFIR P1 Z GPR68 | G protein-coupled receptor 68 | GPR68 |
| BY4741 far1Δ sst2Δ ste2Δ fig1Δ::mTq2 X-2:$P_{TEF1a}$-GPR68-$T_{CYC1b}$ gpa1(468-472)(KIGII>EYNLV) | Kapolka et al., 2020 | DI DCyFIR P1 Q GPR68 | G protein-coupled receptor 68 | GPR68 |
| BY4741 far1Δ sst2Δ ste2Δ fig1Δ::mTq2 X-2:$P_{TEF1a}$-GPR68-$T_{CYC1b}$ gpa1(468-472)(KIGII>EFNLV) | Kapolka et al., 2020 | DI DCyFIR P1 14 GPR68 | G protein-coupled receptor 68 | GPR68 |
| BY4741 far1Δ sst2Δ ste2Δ fig1Δ::mTq2 X-2:$P_{TEF1a}$-GPR68-$T_{CYC1b}$ gpa1(468-472)(KIGII>EINLL) | Kapolka et al., 2020 | DI DCyFIR P1 15 GPR68 | G protein-coupled receptor 68 | GPR68 |
| BY4741 far1Δ sst2Δ ste2Δ fig1Δ::mTq2 X-2:$P_{TEF1a}$-GPR68-$T_{CYC1b}$ gpa1(468-472)(KIGII>DIMLQ) | Kapolka et al., 2020 | DI DCyFIR P1 12 GPR68 | G protein-coupled receptor 68 | GPR68 |
| BY4741 far1Δ sst2Δ ste2Δ fig1Δ::mTq2 X-2:$P_{TEF1a}$-GPR68-$T_{CYC1b}$ gpa1(468-472)(KIGII>QLMLQ) | Kapolka et al., 2020 | DI DCyFIR P1 13 GPR68 | G protein-coupled receptor 68 | GPR68 |
| BY4741 far1Δ sst2Δ ste2Δ fig1Δ::mTq2 X-2:$P_{TEF1a}$-GPR68-$T_{CYC1b}$ gpa1(468-472)(KIGII>QYELL) | Kapolka et al., 2020 | DI DCyFIR P1 S GPR68 | G protein-coupled receptor 68 | GPR68 |

| | | | | |
|---|---|---|---|---|
| BY4741 far1Δ sst2Δ ste2Δ fig1Δ::mTq2 X-2:$P_{TEF1a}$-HCAR2-$T_{CYC1b}$ gpa1(468-472)(KIGII>ECGLY) | Kapolka et al., 2020 | DI DCyFIR P1 I HCAR2 | hydroxycarboxylic acid receptor 2 | HCAR2 |
| BY4741 far1Δ sst2Δ ste2Δ fig1Δ::mTq2 X-2:$P_{TEF1a}$-HCAR2-$T_{CYC1b}$ gpa1(468-472)(KIGII>GCGLY) | Kapolka et al., 2020 | DI DCyFIR P1 O HCAR2 | hydroxycarboxylic acid receptor 2 | HCAR2 |
| BY4741 far1Δ sst2Δ ste2Δ fig1Δ::mTq2 X-2:$P_{TEF1a}$-HCAR2-$T_{CYC1b}$ gpa1(468-472)(KIGII>DCGLF) | Kapolka et al., 2020 | DI DCyFIR P1 T HCAR2 | hydroxycarboxylic acid receptor 2 | HCAR2 |
| BY4741 far1Δ sst2Δ ste2Δ fig1Δ::mTq2 X-2:$P_{TEF1a}$-HCAR2-$T_{CYC1b}$ gpa1(468-472)(KIGII>YIGLC) | Kapolka et al., 2020 | DI DCyFIR P1 Z HCAR2 | hydroxycarboxylic acid receptor 2 | HCAR2 |
| BY4741 far1Δ sst2Δ ste2Δ fig1Δ::mTq2 X-2:$P_{TEF1a}$-HCAR2-$T_{CYC1b}$ gpa1(468-472)(KIGII>EYNLV) | Kapolka et al., 2020 | DI DCyFIR P1 Q HCAR2 | hydroxycarboxylic acid receptor 2 | HCAR2 |
| BY4741 far1Δ sst2Δ ste2Δ fig1Δ::mTq2 X-2:$P_{TEF1a}$-HCAR2-$T_{CYC1b}$ gpa1(468-472)(KIGII>EFNLV) | Kapolka et al., 2020 | DI DCyFIR P1 14 HCAR2 | hydroxycarboxylic acid receptor 2 | HCAR2 |
| BY4741 far1Δ sst2Δ ste2Δ fig1Δ::mTq2 X-2:$P_{TEF1a}$-HCAR2-$T_{CYC1b}$ gpa1(468-472)(KIGII>EINLL) | Kapolka et al., 2020 | DI DCyFIR P1 15 HCAR2 | hydroxycarboxylic acid receptor 2 | HCAR2 |
| BY4741 far1Δ sst2Δ ste2Δ fig1Δ::mTq2 X-2:$P_{TEF1a}$-HCAR2-$T_{CYC1b}$ gpa1(468-472)(KIGII>DIMLQ) | Kapolka et al., 2020 | DI DCyFIR P1 12 HCAR2 | hydroxycarboxylic acid receptor 2 | HCAR2 |
| BY4741 far1Δ sst2Δ ste2Δ fig1Δ::mTq2 X-2:$P_{TEF1a}$-HCAR2-$T_{CYC1b}$ gpa1(468-472)(KIGII>QLMLQ) | Kapolka et al., 2020 | DI DCyFIR P1 13 HCAR2 | hydroxycarboxylic acid receptor 2 | HCAR2 |
| BY4741 far1Δ sst2Δ ste2Δ fig1Δ::mTq2 X-2:$P_{TEF1a}$-HCAR2-$T_{CYC1b}$ gpa1(468-472)(KIGII>QYELL) | Kapolka et al., 2020 | DI DCyFIR P1 S HCAR2 | hydroxycarboxylic acid receptor 2 | HCAR2 |
| BY4741 far1Δ sst2Δ ste2Δ fig1Δ::mTq2 X-2:$P_{TEF1a}$-HCAR3-$T_{CYC1b}$ gpa1(468-472)(KIGII>ECGLY) | Kapolka et al., 2020 | DI DCyFIR P1 I HCAR3 | hydroxycarboxylic acid receptor 3 | HCAR3 |
| BY4741 far1Δ sst2Δ ste2Δ fig1Δ::mTq2 X-2:$P_{TEF1a}$-HCAR3-$T_{CYC1b}$ gpa1(468-472)(KIGII>GCGLY) | Kapolka et al., 2020 | DI DCyFIR P1 O HCAR3 | hydroxycarboxylic acid receptor 3 | HCAR3 |
| BY4741 far1Δ sst2Δ ste2Δ fig1Δ::mTq2 X-2:$P_{TEF1a}$-HCAR3-$T_{CYC1b}$ gpa1(468-472)(KIGII>DCGLF) | Kapolka et al., 2020 | DI DCyFIR P1 T HCAR3 | hydroxycarboxylic acid receptor 3 | HCAR3 |
| BY4741 far1Δ sst2Δ ste2Δ fig1Δ::mTq2 X-2:$P_{TEF1a}$-HCAR3-$T_{CYC1b}$ gpa1(468-472)(KIGII>YIGLC) | Kapolka et al., 2020 | DI DCyFIR P1 Z HCAR3 | hydroxycarboxylic acid receptor 3 | HCAR3 |

| | | | | |
|---|---|---|---|---|
| BY4741 far1Δ sst2Δ ste2Δ fig1Δ::mTq2 X-2:$P_{TEF1a}$-HCAR3-$T_{CYC1b}$ gpa1(468-472)(KIGII>EYNLV) | Kapolka et al., 2020 | DI DCyFIR P1 Q HCAR3 | hydroxycarboxylic acid receptor 3 | HCAR3 |
| BY4741 far1Δ sst2Δ ste2Δ fig1Δ::mTq2 X-2:$P_{TEF1a}$-HCAR3-$T_{CYC1b}$ gpa1(468-472)(KIGII>EFNLV) | Kapolka et al., 2020 | DI DCyFIR P1 14 HCAR3 | hydroxycarboxylic acid receptor 3 | HCAR3 |
| BY4741 far1Δ sst2Δ ste2Δ fig1Δ::mTq2 X-2:$P_{TEF1a}$-HCAR3-$T_{CYC1b}$ gpa1(468-472)(KIGII>EINLL) | Kapolka et al., 2020 | DI DCyFIR P1 15 HCAR3 | hydroxycarboxylic acid receptor 3 | HCAR3 |
| BY4741 far1Δ sst2Δ ste2Δ fig1Δ::mTq2 X-2:$P_{TEF1a}$-HCAR3-$T_{CYC1b}$ gpa1(468-472)(KIGII>DIMLQ) | Kapolka et al., 2020 | DI DCyFIR P1 12 HCAR3 | hydroxycarboxylic acid receptor 3 | HCAR3 |
| BY4741 far1Δ sst2Δ ste2Δ fig1Δ::mTq2 X-2:$P_{TEF1a}$-HCAR3-$T_{CYC1b}$ gpa1(468-472)(KIGII>QLMLQ) | Kapolka et al., 2020 | DI DCyFIR P1 13 HCAR3 | hydroxycarboxylic acid receptor 3 | HCAR3 |
| BY4741 far1Δ sst2Δ ste2Δ fig1Δ::mTq2 X-2:$P_{TEF1a}$-HCAR3-$T_{CYC1b}$ gpa1(468-472)(KIGII>QYELL) | Kapolka et al., 2020 | DI DCyFIR P1 S HCAR3 | hydroxycarboxylic acid receptor 3 | HCAR3 |
| BY4741 far1Δ sst2Δ ste2Δ fig1Δ::mTq2 X-2:$P_{TEF1a}$-HTR4-$T_{CYC1b}$ gpa1(468-472)(KIGII>ECGLY) | Kapolka et al., 2020 | DI DCyFIR P1 I HTR4 | 5-hydroxytryptamine receptor 4 | HTR4 |
| BY4741 far1Δ sst2Δ ste2Δ fig1Δ::mTq2 X-2:$P_{TEF1a}$-HTR4-$T_{CYC1b}$ gpa1(468-472)(KIGII>GCGLY) | Kapolka et al., 2020 | DI DCyFIR P1 O HTR4 | 5-hydroxytryptamine receptor 4 | HTR4 |
| BY4741 far1Δ sst2Δ ste2Δ fig1Δ::mTq2 X-2:$P_{TEF1a}$-HTR4-$T_{CYC1b}$ gpa1(468-472)(KIGII>DCGLF) | Kapolka et al., 2020 | DI DCyFIR P1 T HTR4 | 5-hydroxytryptamine receptor 4 | HTR4 |
| BY4741 far1Δ sst2Δ ste2Δ fig1Δ::mTq2 X-2:$P_{TEF1a}$-HTR4-$T_{CYC1b}$ gpa1(468-472)(KIGII>YIGLC) | Kapolka et al., 2020 | DI DCyFIR P1 Z HTR4 | 5-hydroxytryptamine receptor 4 | HTR4 |
| BY4741 far1Δ sst2Δ ste2Δ fig1Δ::mTq2 X-2:$P_{TEF1a}$-HTR4-$T_{CYC1b}$ gpa1(468-472)(KIGII>EYNLV) | Kapolka et al., 2020 | DI DCyFIR P1 Q HTR4 | 5-hydroxytryptamine receptor 4 | HTR4 |
| BY4741 far1Δ sst2Δ ste2Δ fig1Δ::mTq2 X-2:$P_{TEF1a}$-HTR4-$T_{CYC1b}$ gpa1(468-472)(KIGII>EFNLV) | Kapolka et al., 2020 | DI DCyFIR P1 14 HTR4 | 5-hydroxytryptamine receptor 4 | HTR4 |

| | | | | |
|---|---|---|---|---|
| BY4741 far1Δ sst2Δ ste2Δ fig1Δ::mTq2 X-2:$P_{TEF1a}$-HTR4-$T_{CYC1b}$ gpa1(468-472)(KIGII>EINLL) | Kapolka et al., 2020 | DI DCyFIR P1 15 HTR4 | 5-hydroxytryptamine receptor 4 | HTR4 |
| BY4741 far1Δ sst2Δ ste2Δ fig1Δ::mTq2 X-2:$P_{TEF1a}$-HTR4-$T_{CYC1b}$ gpa1(468-472)(KIGII>DIMLQ) | Kapolka et al., 2020 | DI DCyFIR P1 12 HTR4 | 5-hydroxytryptamine receptor 4 | HTR4 |
| BY4741 far1Δ sst2Δ ste2Δ fig1Δ::mTq2 X-2:$P_{TEF1a}$-HTR4-$T_{CYC1b}$ gpa1(468-472)(KIGII>QLMLQ) | Kapolka et al., 2020 | DI DCyFIR P1 13 HTR4 | 5-hydroxytryptamine receptor 4 | HTR4 |
| BY4741 far1Δ sst2Δ ste2Δ fig1Δ::mTq2 X-2:$P_{TEF1a}$-HTR4-$T_{CYC1b}$ gpa1(468-472)(KIGII>QYELL) | Kapolka et al., 2020 | DI DCyFIR P1 S HTR4 | 5-hydroxytryptamine receptor 4 | HTR4 |
| BY4741 far1Δ sst2Δ ste2Δ fig1Δ::mTq2 X-2:PTEF1a-LPAR3-TCYC1b gpa1(468-472)(KIGII>ECGLY) | This paper | DI DCyFIR P1 I LPAR3 | lysophosphatidic acid receptor 3 | LPAR3 |
| BY4741 far1Δ sst2Δ ste2Δ fig1Δ::mTq2 X-2:PTEF1a-LPAR3-TCYC1b gpa1(468-472)(KIGII>GCGLY) | This paper | DI DCyFIR P1 O LPAR3 | lysophosphatidic acid receptor 3 | LPAR3 |
| BY4741 far1Δ sst2Δ ste2Δ fig1Δ::mTq2 X-2:PTEF1a-LPAR3-TCYC1b gpa1(468-472)(KIGII>DCGLF) | This paper | DI DCyFIR P1 T LPAR3 | lysophosphatidic acid receptor 3 | LPAR3 |
| BY4741 far1Δ sst2Δ ste2Δ fig1Δ::mTq2 X-2:PTEF1a-LPAR3-TCYC1b gpa1(468-472)(KIGII>YIGLC) | This paper | DI DCyFIR P1 Z LPAR3 | lysophosphatidic acid receptor 3 | LPAR3 |
| BY4741 far1Δ sst2Δ ste2Δ fig1Δ::mTq2 X-2:PTEF1a-LPAR3-TCYC1b gpa1(468-472)(KIGII>EYNLV) | This paper | DI DCyFIR P1 Q LPAR3 | lysophosphatidic acid receptor 3 | LPAR3 |
| BY4741 far1Δ sst2Δ ste2Δ fig1Δ::mTq2 X-2:PTEF1a-LPAR3-TCYC1b gpa1(468-472)(KIGII>EFNLV) | This paper | DI DCyFIR P1 14 LPAR3 | lysophosphatidic acid receptor 3 | LPAR3 |
| BY4741 far1Δ sst2Δ ste2Δ fig1Δ::mTq2 X-2:PTEF1a-LPAR3-TCYC1b gpa1(468-472)(KIGII>EINLL) | This paper | DI DCyFIR P1 15 LPAR3 | lysophosphatidic acid receptor 3 | LPAR3 |
| BY4741 far1Δ sst2Δ ste2Δ fig1Δ::mTq2 X-2:PTEF1a-LPAR3-TCYC1b gpa1(468-472)(KIGII>DIMLQ) | This paper | DI DCyFIR P1 12 LPAR3 | lysophosphatidic acid receptor 3 | LPAR3 |
| BY4741 far1Δ sst2Δ ste2Δ fig1Δ::mTq2 X-2:PTEF1a-LPAR3-TCYC1b gpa1(468-472)(KIGII>QLMLQ) | This paper | DI DCyFIR P1 13 LPAR3 | lysophosphatidic acid receptor 3 | LPAR3 |

| | | | | |
|---|---|---|---|---|
| BY4741 far1Δ sst2Δ ste2Δ fig1Δ::mTq2 X-2:PTEF1a-LPAR3-TCYC1b gpa1(468-472)(KIGII>QYELL) | This paper | DI DCyFIR P1 S LPAR3 | lysophosphatidic acid receptor 3 | LPAR3 |
| BY4741 far1Δ sst2Δ ste2Δ fig1Δ::mTq2 X-2:$P_{TEF1a}$-LPAR4-$T_{CYC1b}$ gpa1(468-472)(KIGII>ECGLY) | Kapolka et al., 2020 | DI DCyFIR P1 I LPAR4 | lysophosphatidic acid receptor 4 | LPAR4 |
| BY4741 far1Δ sst2Δ ste2Δ fig1Δ::mTq2 X-2:$P_{TEF1a}$-LPAR4-$T_{CYC1b}$ gpa1(468-472)(KIGII>GCGLY) | Kapolka et al., 2020 | DI DCyFIR P1 O LPAR4 | lysophosphatidic acid receptor 4 | LPAR4 |
| BY4741 far1Δ sst2Δ ste2Δ fig1Δ::mTq2 X-2:$P_{TEF1a}$-LPAR4-$T_{CYC1b}$ gpa1(468-472)(KIGII>DCGLF) | Kapolka et al., 2020 | DI DCyFIR P1 T LPAR4 | lysophosphatidic acid receptor 4 | LPAR4 |
| BY4741 far1Δ sst2Δ ste2Δ fig1Δ::mTq2 X-2:$P_{TEF1a}$-LPAR4-$T_{CYC1b}$ gpa1(468-472)(KIGII>YIGLC) | Kapolka et al., 2020 | DI DCyFIR P1 Z LPAR4 | lysophosphatidic acid receptor 4 | LPAR4 |
| BY4741 far1Δ sst2Δ ste2Δ fig1Δ::mTq2 X-2:$P_{TEF1a}$-LPAR4-$T_{CYC1b}$ gpa1(468-472)(KIGII>EYNLV) | Kapolka et al., 2020 | DI DCyFIR P1 Q LPAR4 | lysophosphatidic acid receptor 4 | LPAR4 |
| BY4741 far1Δ sst2Δ ste2Δ fig1Δ::mTq2 X-2:$P_{TEF1a}$-LPAR4-$T_{CYC1b}$ gpa1(468-472)(KIGII>EFNLV) | Kapolka et al., 2020 | DI DCyFIR P1 14 LPAR4 | lysophosphatidic acid receptor 4 | LPAR4 |
| BY4741 far1Δ sst2Δ ste2Δ fig1Δ::mTq2 X-2:$P_{TEF1a}$-LPAR4-$T_{CYC1b}$ gpa1(468-472)(KIGII>EINLL) | Kapolka et al., 2020 | DI DCyFIR P1 15 LPAR4 | lysophosphatidic acid receptor 4 | LPAR4 |
| BY4741 far1Δ sst2Δ ste2Δ fig1Δ::mTq2 X-2:$P_{TEF1a}$-LPAR4-$T_{CYC1b}$ gpa1(468-472)(KIGII>DIMLQ) | Kapolka et al., 2020 | DI DCyFIR P1 12 LPAR4 | lysophosphatidic acid receptor 4 | LPAR4 |
| BY4741 far1Δ sst2Δ ste2Δ fig1Δ::mTq2 X-2:$P_{TEF1a}$-LPAR4-$T_{CYC1b}$ gpa1(468-472)(KIGII>QLMLQ) | Kapolka et al., 2020 | DI DCyFIR P1 13 LPAR4 | lysophosphatidic acid receptor 4 | LPAR4 |
| BY4741 far1Δ sst2Δ ste2Δ fig1Δ::mTq2 X-2:$P_{TEF1a}$-LPAR4-$T_{CYC1b}$ gpa1(468-472)(KIGII>QYELL) | Kapolka et al., 2020 | DI DCyFIR P1 S LPAR4 | lysophosphatidic acid receptor 4 | LPAR4 |
| BY4741 far1Δ sst2Δ ste2Δ fig1Δ::mTq2 X-2:$P_{TEF1a}$-MTNR1A-$T_{CYC1b}$ gpa1(468-472)(KIGII>ECGLY) | Kapolka et al., 2020 | DI DCyFIR P1 I MTNR1A | melatonin receptor 1A | MTNR1A |
| BY4741 far1Δ sst2Δ ste2Δ fig1Δ::mTq2 X-2:$P_{TEF1a}$-MTNR1A-$T_{CYC1b}$ gpa1(468-472)(KIGII>GCGLY) | Kapolka et al., 2020 | DI DCyFIR P1 O MTNR1A | melatonin receptor 1A | MTNR1A |
| BY4741 far1Δ sst2Δ ste2Δ fig1Δ::mTq2 X-2:$P_{TEF1a}$-MTNR1A-$T_{CYC1b}$ gpa1(468-472)(KIGII>DCGLF) | Kapolka et al., 2020 | DI DCyFIR P1 T MTNR1A | melatonin receptor 1A | MTNR1A |

| | | | | |
|---|---|---|---|---|
| BY4741 far1Δ sst2Δ ste2Δ fig1Δ::mTq2 X-2:$P_{TEF1a}$-MTNR1A-$T_{CYC1b}$ gpa1(468-472)(KIGII>YIGLC) | Kapolka et al., 2020 | DI DCyFIR P1 Z MTNR1A | melatonin receptor 1A | MTNR1A |
| BY4741 far1Δ sst2Δ ste2Δ fig1Δ::mTq2 X-2:$P_{TEF1a}$-MTNR1A-$T_{CYC1b}$ gpa1(468-472)(KIGII>EYNLV) | Kapolka et al., 2020 | DI DCyFIR P1 Q MTNR1A | melatonin receptor 1A | MTNR1A |
| BY4741 far1Δ sst2Δ ste2Δ fig1Δ::mTq2 X-2:$P_{TEF1a}$-MTNR1A-$T_{CYC1b}$ gpa1(468-472)(KIGII>EFNLV) | Kapolka et al., 2020 | DI DCyFIR P1 14 MTNR1A | melatonin receptor 1A | MTNR1A |
| BY4741 far1Δ sst2Δ ste2Δ fig1Δ::mTq2 X-2:$P_{TEF1a}$-MTNR1A-$T_{CYC1b}$ gpa1(468-472)(KIGII>EINLL) | Kapolka et al., 2020 | DI DCyFIR P1 15 MTNR1A | melatonin receptor 1A | MTNR1A |
| BY4741 far1Δ sst2Δ ste2Δ fig1Δ::mTq2 X-2:$P_{TEF1a}$-MTNR1A-$T_{CYC1b}$ gpa1(468-472)(KIGII>DIMLQ) | Kapolka et al., 2020 | DI DCyFIR P1 12 MTNR1A | melatonin receptor 1A | MTNR1A |
| BY4741 far1Δ sst2Δ ste2Δ fig1Δ::mTq2 X-2:$P_{TEF1a}$-MTNR1A-$T_{CYC1b}$ gpa1(468-472)(KIGII>QLMLQ) | Kapolka et al., 2020 | DI DCyFIR P1 13 MTNR1A | melatonin receptor 1A | MTNR1A |
| BY4741 far1Δ sst2Δ ste2Δ fig1Δ::mTq2 X-2:$P_{TEF1a}$-MTNR1A-$T_{CYC1b}$ gpa1(468-472)(KIGII>QYELL) | Kapolka et al., 2020 | DI DCyFIR P1 S MTNR1A | melatonin receptor 1A | MTNR1A |
| BY4741 far1Δ sst2Δ ste2Δ fig1Δ::mTq2 X-2:$P_{TEF1a}$-MTNR1B-$T_{CYC1b}$ gpa1(468-472)(KIGII>ECGLY) | Kapolka et al., 2020 | DI DCyFIR P1 I MTNR1B | melatonin receptor 1B | MTNR1B |
| BY4741 far1Δ sst2Δ ste2Δ fig1Δ::mTq2 X-2:$P_{TEF1a}$-MTNR1B-$T_{CYC1b}$ gpa1(468-472)(KIGII>GCGLY) | Kapolka et al., 2020 | DI DCyFIR P1 O MTNR1B | melatonin receptor 1B | MTNR1B |
| BY4741 far1Δ sst2Δ ste2Δ fig1Δ::mTq2 X-2:$P_{TEF1a}$-MTNR1B-$T_{CYC1b}$ gpa1(468-472)(KIGII>DCGLF) | Kapolka et al., 2020 | DI DCyFIR P1 T MTNR1B | melatonin receptor 1B | MTNR1B |
| BY4741 far1Δ sst2Δ ste2Δ fig1Δ::mTq2 X-2:$P_{TEF1a}$-MTNR1B-$T_{CYC1b}$ gpa1(468-472)(KIGII>YIGLC) | Kapolka et al., 2020 | DI DCyFIR P1 Z MTNR1B | melatonin receptor 1B | MTNR1B |
| BY4741 far1Δ sst2Δ ste2Δ fig1Δ::mTq2 X-2:$P_{TEF1a}$-MTNR1B-$T_{CYC1b}$ gpa1(468-472)(KIGII>EYNLV) | Kapolka et al., 2020 | DI DCyFIR P1 Q MTNR1B | melatonin receptor 1B | MTNR1B |
| BY4741 far1Δ sst2Δ ste2Δ fig1Δ::mTq2 X-2:$P_{TEF1a}$-MTNR1B-$T_{CYC1b}$ gpa1(468-472)(KIGII>EFNLV) | Kapolka et al., 2020 | DI DCyFIR P1 14 MTNR1B | melatonin receptor 1B | MTNR1B |
| BY4741 far1Δ sst2Δ ste2Δ fig1Δ::mTq2 X-2:$P_{TEF1a}$-MTNR1B-$T_{CYC1b}$ gpa1(468-472)(KIGII>EINLL) | Kapolka et al., 2020 | DI DCyFIR P1 15 MTNR1B | melatonin receptor 1B | MTNR1B |

| | | | | |
|---|---|---|---|---|
| BY4741 far1Δ sst2Δ ste2Δ fig1Δ::mTq2 X-2:$P_{TEF1a}$-MTNR1B-$T_{CYC1b}$ gpa1(468-472)(KIGII>DIMLQ) | Kapolka et al., 2020 | DI DCyFIR P1 12 MTNR1B | melatonin receptor 1B | MTNR1B |
| BY4741 far1Δ sst2Δ ste2Δ fig1Δ::mTq2 X-2:$P_{TEF1a}$-MTNR1B-$T_{CYC1b}$ gpa1(468-472)(KIGII>QLMLQ) | Kapolka et al., 2020 | DI DCyFIR P1 13 MTNR1B | melatonin receptor 1B | MTNR1B |
| BY4741 far1Δ sst2Δ ste2Δ fig1Δ::mTq2 X-2:$P_{TEF1a}$-MTNR1B-$T_{CYC1b}$ gpa1(468-472)(KIGII>QYELL) | Kapolka et al., 2020 | DI DCyFIR P1 S MTNR1B | melatonin receptor 1B | MTNR1B |
| BY4741 far1Δ sst2Δ ste2Δ fig1Δ::mTq2 X-2:$P_{TEF1a}$-PTAFR-$T_{CYC1b}$ gpa1(468-472)(KIGII>ECGLY) | Kapolka et al., 2020 | DI DCyFIR P1 I PTAFR | platelet-activating factor receptor | PTAFR |
| BY4741 far1Δ sst2Δ ste2Δ fig1Δ::mTq2 X-2:$P_{TEF1a}$-PTAFR-$T_{CYC1b}$ gpa1(468-472)(KIGII>GCGLY) | Kapolka et al., 2020 | DI DCyFIR P1 O PTAFR | platelet-activating factor receptor | PTAFR |
| BY4741 far1Δ sst2Δ ste2Δ fig1Δ::mTq2 X-2:$P_{TEF1a}$-PTAFR-$T_{CYC1b}$ gpa1(468-472)(KIGII>DCGLF) | Kapolka et al., 2020 | DI DCyFIR P1 T PTAFR | platelet-activating factor receptor | PTAFR |
| BY4741 far1Δ sst2Δ ste2Δ fig1Δ::mTq2 X-2:$P_{TEF1a}$-PTAFR-$T_{CYC1b}$ gpa1(468-472)(KIGII>YIGLC) | Kapolka et al., 2020 | DI DCyFIR P1 Z PTAFR | platelet-activating factor receptor | PTAFR |
| BY4741 far1Δ sst2Δ ste2Δ fig1Δ::mTq2 X-2:$P_{TEF1a}$-PTAFR-$T_{CYC1b}$ gpa1(468-472)(KIGII>EYNLV) | Kapolka et al., 2020 | DI DCyFIR P1 Q PTAFR | platelet-activating factor receptor | PTAFR |
| BY4741 far1Δ sst2Δ ste2Δ fig1Δ::mTq2 X-2:$P_{TEF1a}$-PTAFR-$T_{CYC1b}$ gpa1(468-472)(KIGII>EFNLV) | Kapolka et al., 2020 | DI DCyFIR P1 14 PTAFR | platelet-activating factor receptor | PTAFR |
| BY4741 far1Δ sst2Δ ste2Δ fig1Δ::mTq2 X-2:$P_{TEF1a}$-PTAFR-$T_{CYC1b}$ gpa1(468-472)(KIGII>EINLL) | Kapolka et al., 2020 | DI DCyFIR P1 15 PTAFR | platelet-activating factor receptor | PTAFR |
| BY4741 far1Δ sst2Δ ste2Δ fig1Δ::mTq2 X-2:$P_{TEF1a}$-PTAFR-$T_{CYC1b}$ gpa1(468-472)(KIGII>DIMLQ) | Kapolka et al., 2020 | DI DCyFIR P1 12 PTAFR | platelet-activating factor receptor | PTAFR |
| BY4741 far1Δ sst2Δ ste2Δ fig1Δ::mTq2 X-2:$P_{TEF1a}$-PTAFR-$T_{CYC1b}$ gpa1(468-472)(KIGII>QLMLQ) | Kapolka et al., 2020 | DI DCyFIR P1 13 PTAFR | platelet-activating | PTAFR |

| | | | | |
|---|---|---|---|---|
| | | | factor receptor | |
| BY4741 far1Δ sst2Δ ste2Δ fig1Δ::mTq2 X-2:$P_{TEF1a}$-PTAFR-$T_{CYC1b}$ gpa1(468-472)(KIGII>QYELL) | Kapolka et al., 2020 | DI DCyFIR P1 S PTAFR | platelet-activating factor receptor | PTAFR |
| BY4741 far1Δ sst2Δ ste2Δ fig1Δ::mTq2 X-2:$P_{TEF1a}$-PTGER3-$T_{CYC1b}$ gpa1(468-472)(KIGII>ECGLY) | Kapolka et al., 2020 | DI DCyFIR P1 I PTGER3 | prostaglandin EP3 receptor | PTGER3 |
| BY4741 far1Δ sst2Δ ste2Δ fig1Δ::mTq2 X-2:$P_{TEF1a}$-PTGER3-$T_{CYC1b}$ gpa1(468-472)(KIGII>GCGLY) | Kapolka et al., 2020 | DI DCyFIR P1 O PTGER3 | prostaglandin EP3 receptor | PTGER3 |
| BY4741 far1Δ sst2Δ ste2Δ fig1Δ::mTq2 X-2:$P_{TEF1a}$-PTGER3-$T_{CYC1b}$ gpa1(468-472)(KIGII>DCGLF) | Kapolka et al., 2020 | DI DCyFIR P1 T PTGER3 | prostaglandin EP3 receptor | PTGER3 |
| BY4741 far1Δ sst2Δ ste2Δ fig1Δ::mTq2 X-2:$P_{TEF1a}$-PTGER3-$T_{CYC1b}$ gpa1(468-472)(KIGII>YIGLC) | Kapolka et al., 2020 | DI DCyFIR P1 Z PTGER3 | prostaglandin EP3 receptor | PTGER3 |
| BY4741 far1Δ sst2Δ ste2Δ fig1Δ::mTq2 X-2:$P_{TEF1a}$-PTGER3-$T_{CYC1b}$ gpa1(468-472)(KIGII>EYNLV) | Kapolka et al., 2020 | DI DCyFIR P1 Q PTGER3 | prostaglandin EP3 receptor | PTGER3 |
| BY4741 far1Δ sst2Δ ste2Δ fig1Δ::mTq2 X-2:$P_{TEF1a}$-PTGER3-$T_{CYC1b}$ gpa1(468-472)(KIGII>EFNLV) | Kapolka et al., 2020 | DI DCyFIR P1 14 PTGER3 | prostaglandin EP3 receptor | PTGER3 |
| BY4741 far1Δ sst2Δ ste2Δ fig1Δ::mTq2 X-2:$P_{TEF1a}$-PTGER3-$T_{CYC1b}$ gpa1(468-472)(KIGII>EINLL) | Kapolka et al., 2020 | DI DCyFIR P1 15 PTGER3 | prostaglandin EP3 receptor | PTGER3 |
| BY4741 far1Δ sst2Δ ste2Δ fig1Δ::mTq2 X-2:$P_{TEF1a}$-PTGER3-$T_{CYC1b}$ gpa1(468-472)(KIGII>DIMLQ) | Kapolka et al., 2020 | DI DCyFIR P1 12 PTGER3 | prostaglandin EP3 receptor | PTGER3 |
| BY4741 far1Δ sst2Δ ste2Δ fig1Δ::mTq2 X-2:$P_{TEF1a}$-PTGER3-$T_{CYC1b}$ gpa1(468-472)(KIGII>QLMLQ) | Kapolka et al., 2020 | DI DCyFIR P1 13 PTGER3 | prostaglandin EP3 receptor | PTGER3 |
| BY4741 far1Δ sst2Δ ste2Δ fig1Δ::mTq2 X-2:$P_{TEF1a}$-PTGER3-$T_{CYC1b}$ gpa1(468-472)(KIGII>QYELL) | Kapolka et al., 2020 | DI DCyFIR P1 S PTGER3 | prostaglandin EP3 receptor | PTGER3 |
| BY4741 far1Δ sst2Δ ste2Δ fig1Δ::mTq2 X-2:$P_{TEF1a}$-S1PR1-$T_{CYC1b}$ gpa1(468-472)(KIGII>ECGLY) | Kapolka et al., 2020 | DI DCyFIR P1 I S1PR1 | sphingosine-1-phosphate receptor 1 | S1PR1 |
| BY4741 far1Δ sst2Δ ste2Δ fig1Δ::mTq2 X-2:$P_{TEF1a}$-S1PR1-$T_{CYC1b}$ gpa1(468-472)(KIGII>GCGLY) | Kapolka et al., 2020 | DI DCyFIR P1 O S1PR1 | sphingosine-1-phosphate receptor 1 | S1PR1 |

| | | | | |
|---|---|---|---|---|
| BY4741 far1Δ sst2Δ ste2Δ fig1Δ::mTq2 X-2:$P_{TEF1a}$-S1PR1-$T_{CYC1b}$ gpa1(468-472)(KIGII>DCGLF) | Kapolka et al., 2020 | DI DCyFIR P1 T S1PR1 | sphingosine-1-phosphate receptor 1 | S1PR1 |
| BY4741 far1Δ sst2Δ ste2Δ fig1Δ::mTq2 X-2:$P_{TEF1a}$-S1PR1-$T_{CYC1b}$ gpa1(468-472)(KIGII>YIGLC) | Kapolka et al., 2020 | DI DCyFIR P1 Z S1PR1 | sphingosine-1-phosphate receptor 1 | S1PR1 |
| BY4741 far1Δ sst2Δ ste2Δ fig1Δ::mTq2 X-2:$P_{TEF1a}$-S1PR1-$T_{CYC1b}$ gpa1(468-472)(KIGII>EYNLV) | Kapolka et al., 2020 | DI DCyFIR P1 Q S1PR1 | sphingosine-1-phosphate receptor 1 | S1PR1 |
| BY4741 far1Δ sst2Δ ste2Δ fig1Δ::mTq2 X-2:$P_{TEF1a}$-S1PR1-$T_{CYC1b}$ gpa1(468-472)(KIGII>EFNLV) | Kapolka et al., 2020 | DI DCyFIR P1 14 S1PR1 | sphingosine-1-phosphate receptor 1 | S1PR1 |
| BY4741 far1Δ sst2Δ ste2Δ fig1Δ::mTq2 X-2:$P_{TEF1a}$-S1PR1-$T_{CYC1b}$ gpa1(468-472)(KIGII>EINLL) | Kapolka et al., 2020 | DI DCyFIR P1 15 S1PR1 | sphingosine-1-phosphate receptor 1 | S1PR1 |
| BY4741 far1Δ sst2Δ ste2Δ fig1Δ::mTq2 X-2:$P_{TEF1a}$-S1PR1-$T_{CYC1b}$ gpa1(468-472)(KIGII>DIMLQ) | Kapolka et al., 2020 | DI DCyFIR P1 12 S1PR1 | sphingosine-1-phosphate receptor 1 | S1PR1 |
| BY4741 far1Δ sst2Δ ste2Δ fig1Δ::mTq2 X-2:$P_{TEF1a}$-S1PR1-$T_{CYC1b}$ gpa1(468-472)(KIGII>QLMLQ) | Kapolka et al., 2020 | DI DCyFIR P1 13 S1PR1 | sphingosine-1-phosphate receptor 1 | S1PR1 |
| BY4741 far1Δ sst2Δ ste2Δ fig1Δ::mTq2 X-2:$P_{TEF1a}$-S1PR1-$T_{CYC1b}$ gpa1(468-472)(KIGII>QYELL) | Kapolka et al., 2020 | DI DCyFIR P1 S S1PR1 | sphingosine-1-phosphate receptor 1 | S1PR1 |
| BY4741 far1Δ sst2Δ ste2Δ fig1Δ::mTq2 X-2:$P_{TEF1a}$-S1PR2-$T_{CYC1b}$ gpa1(468-472)(KIGII>ECGLY) | Kapolka et al., 2020 | DI DCyFIR P1 I S1PR2 | sphingosine-1-phosphate receptor 2 | S1PR2 |
| BY4741 far1Δ sst2Δ ste2Δ fig1Δ::mTq2 X-2:$P_{TEF1a}$-S1PR2-$T_{CYC1b}$ gpa1(468-472)(KIGII>GCGLY) | Kapolka et al., 2020 | DI DCyFIR P1 O S1PR2 | sphingosine-1-phosphate receptor 2 | S1PR2 |
| BY4741 far1Δ sst2Δ ste2Δ fig1Δ::mTq2 X-2:$P_{TEF1a}$-S1PR2-$T_{CYC1b}$ gpa1(468-472)(KIGII>DCGLF) | Kapolka et al., 2020 | DI DCyFIR P1 T S1PR2 | sphingosine-1-phosphate receptor 2 | S1PR2 |
| BY4741 far1Δ sst2Δ ste2Δ fig1Δ::mTq2 X-2:$P_{TEF1a}$-S1PR2-$T_{CYC1b}$ gpa1(468-472)(KIGII>YIGLC) | Kapolka et al., 2020 | DI DCyFIR P1 Z S1PR2 | sphingosine-1-phosphate receptor 2 | S1PR2 |
| BY4741 far1Δ sst2Δ ste2Δ fig1Δ::mTq2 X-2:$P_{TEF1a}$-S1PR2-$T_{CYC1b}$ gpa1(468-472)(KIGII>EYNLV) | Kapolka et al., 2020 | DI DCyFIR P1 Q S1PR2 | sphingosine-1-phosphate receptor 2 | S1PR2 |
| BY4741 far1Δ sst2Δ ste2Δ fig1Δ::mTq2 X-2:$P_{TEF1a}$-S1PR2-$T_{CYC1b}$ gpa1(468-472)(KIGII>EFNLV) | Kapolka et al., 2020 | DI DCyFIR P1 14 S1PR2 | sphingosine-1-phosphate receptor 2 | S1PR2 |

| | | | | |
|---|---|---|---|---|
| BY4741 far1Δ sst2Δ ste2Δ fig1Δ::mTq2 X-2:$P_{TEF1a}$-S1PR2-$T_{CYC1b}$ gpa1(468-472)(KIGII>EINLL) | Kapolka et al., 2020 | DI DCyFIR P1 15 S1PR2 | sphingosine-1-phosphate receptor 2 | S1PR2 |
| BY4741 far1Δ sst2Δ ste2Δ fig1Δ::mTq2 X-2:$P_{TEF1a}$-S1PR2-$T_{CYC1b}$ gpa1(468-472)(KIGII>DIMLQ) | Kapolka et al., 2020 | DI DCyFIR P1 12 S1PR2 | sphingosine-1-phosphate receptor 2 | S1PR2 |
| BY4741 far1Δ sst2Δ ste2Δ fig1Δ::mTq2 X-2:$P_{TEF1a}$-S1PR2-$T_{CYC1b}$ gpa1(468-472)(KIGII>QLMLQ) | Kapolka et al., 2020 | DI DCyFIR P1 13 S1PR2 | sphingosine-1-phosphate receptor 2 | S1PR2 |
| BY4741 far1Δ sst2Δ ste2Δ fig1Δ::mTq2 X-2:$P_{TEF1a}$-S1PR2-$T_{CYC1b}$ gpa1(468-472)(KIGII>QYELL) | Kapolka et al., 2020 | DI DCyFIR P1 S S1PR2 | sphingosine-1-phosphate receptor 2 | S1PR2 |
| BY4741 far1Δ sst2Δ ste2Δ fig1Δ::mTq2 X-2:$P_{TEF1a}$-S1PR3-$T_{CYC1b}$ gpa1(468-472)(KIGII>ECGLY) | Kapolka et al., 2020 | DI DCyFIR P1 I S1PR3 | sphingosine-1-phosphate receptor 3 | S1PR3 |
| BY4741 far1Δ sst2Δ ste2Δ fig1Δ::mTq2 X-2:$P_{TEF1a}$-S1PR3-$T_{CYC1b}$ gpa1(468-472)(KIGII>GCGLY) | Kapolka et al., 2020 | DI DCyFIR P1 O S1PR3 | sphingosine-1-phosphate receptor 3 | S1PR3 |
| BY4741 far1Δ sst2Δ ste2Δ fig1Δ::mTq2 X-2:$P_{TEF1a}$-S1PR3-$T_{CYC1b}$ gpa1(468-472)(KIGII>DCGLF) | Kapolka et al., 2020 | DI DCyFIR P1 T S1PR3 | sphingosine-1-phosphate receptor 3 | S1PR3 |
| BY4741 far1Δ sst2Δ ste2Δ fig1Δ::mTq2 X-2:$P_{TEF1a}$-S1PR3-$T_{CYC1b}$ gpa1(468-472)(KIGII>YIGLC) | Kapolka et al., 2020 | DI DCyFIR P1 Z S1PR3 | sphingosine-1-phosphate receptor 3 | S1PR3 |
| BY4741 far1Δ sst2Δ ste2Δ fig1Δ::mTq2 X-2:$P_{TEF1a}$-S1PR3-$T_{CYC1b}$ gpa1(468-472)(KIGII>EYNLV) | Kapolka et al., 2020 | DI DCyFIR P1 Q S1PR3 | sphingosine-1-phosphate receptor 3 | S1PR3 |
| BY4741 far1Δ sst2Δ ste2Δ fig1Δ::mTq2 X-2:$P_{TEF1a}$-S1PR3-$T_{CYC1b}$ gpa1(468-472)(KIGII>EFNLV) | Kapolka et al., 2020 | DI DCyFIR P1 14 S1PR3 | sphingosine-1-phosphate receptor 3 | S1PR3 |
| BY4741 far1Δ sst2Δ ste2Δ fig1Δ::mTq2 X-2:$P_{TEF1a}$-S1PR3-$T_{CYC1b}$ gpa1(468-472)(KIGII>EINLL) | Kapolka et al., 2020 | DI DCyFIR P1 15 S1PR3 | sphingosine-1-phosphate receptor 3 | S1PR3 |
| BY4741 far1Δ sst2Δ ste2Δ fig1Δ::mTq2 X-2:$P_{TEF1a}$-S1PR3-$T_{CYC1b}$ gpa1(468-472)(KIGII>DIMLQ) | Kapolka et al., 2020 | DI DCyFIR P1 12 S1PR3 | sphingosine-1-phosphate receptor 3 | S1PR3 |
| BY4741 far1Δ sst2Δ ste2Δ fig1Δ::mTq2 X-2:$P_{TEF1a}$-S1PR3-$T_{CYC1b}$ gpa1(468-472)(KIGII>QLMLQ) | Kapolka et al., 2020 | DI DCyFIR P1 13 S1PR3 | sphingosine-1-phosphate receptor 3 | S1PR3 |
| BY4741 far1Δ sst2Δ ste2Δ fig1Δ::mTq2 X-2:$P_{TEF1a}$-S1PR3-$T_{CYC1b}$ gpa1(468-472)(KIGII>QYELL) | Kapolka et al., 2020 | DI DCyFIR P1 S S1PR3 | sphingosine-1-phosphate receptor 3 | S1PR3 |

| | | | | |
|---|---|---|---|---|
| BY4741 far1Δ sst2Δ ste2Δ fig1Δ::mTq2 X-2:$P_{TEF1a}$-SSTR5-$T_{CYC1b}$ gpa1(468-472)(KIGII>ECGLY) | Kapolka et al., 2020 | DI DCyFIR P1 I SSTR5 | somatostatin receptor type 5 | SSTR5 |
| BY4741 far1Δ sst2Δ ste2Δ fig1Δ::mTq2 X-2:$P_{TEF1a}$-SSTR5-$T_{CYC1b}$ gpa1(468-472)(KIGII>GCGLY) | Kapolka et al., 2020 | DI DCyFIR P1 O SSTR5 | somatostatin receptor type 5 | SSTR5 |
| BY4741 far1Δ sst2Δ ste2Δ fig1Δ::mTq2 X-2:$P_{TEF1a}$-SSTR5-$T_{CYC1b}$ gpa1(468-472)(KIGII>DCGLF) | Kapolka et al., 2020 | DI DCyFIR P1 T SSTR5 | somatostatin receptor type 5 | SSTR5 |
| BY4741 far1Δ sst2Δ ste2Δ fig1Δ::mTq2 X-2:$P_{TEF1a}$-SSTR5-$T_{CYC1b}$ gpa1(468-472)(KIGII>YIGLC) | Kapolka et al., 2020 | DI DCyFIR P1 Z SSTR5 | somatostatin receptor type 5 | SSTR5 |
| BY4741 far1Δ sst2Δ ste2Δ fig1Δ::mTq2 X-2:$P_{TEF1a}$-SSTR5-$T_{CYC1b}$ gpa1(468-472)(KIGII>EYNLV) | Kapolka et al., 2020 | DI DCyFIR P1 Q SSTR5 | somatostatin receptor type 5 | SSTR5 |
| BY4741 far1Δ sst2Δ ste2Δ fig1Δ::mTq2 X-2:$P_{TEF1a}$-SSTR5-$T_{CYC1b}$ gpa1(468-472)(KIGII>EFNLV) | Kapolka et al., 2020 | DI DCyFIR P1 14 SSTR5 | somatostatin receptor type 5 | SSTR5 |
| BY4741 far1Δ sst2Δ ste2Δ fig1Δ::mTq2 X-2:$P_{TEF1a}$-SSTR5-$T_{CYC1b}$ gpa1(468-472)(KIGII>EINLL) | Kapolka et al., 2020 | DI DCyFIR P1 15 SSTR5 | somatostatin receptor type 5 | SSTR5 |
| BY4741 far1Δ sst2Δ ste2Δ fig1Δ::mTq2 X-2:$P_{TEF1a}$-SSTR5-$T_{CYC1b}$ gpa1(468-472)(KIGII>DIMLQ) | Kapolka et al., 2020 | DI DCyFIR P1 12 SSTR5 | somatostatin receptor type 5 | SSTR5 |
| BY4741 far1Δ sst2Δ ste2Δ fig1Δ::mTq2 X-2:$P_{TEF1a}$-SSTR5-$T_{CYC1b}$ gpa1(468-472)(KIGII>QLMLQ) | Kapolka et al., 2020 | DI DCyFIR P1 13 SSTR5 | somatostatin receptor type 5 | SSTR5 |
| BY4741 far1Δ sst2Δ ste2Δ fig1Δ::mTq2 X-2:$P_{TEF1a}$-SSTR5-$T_{CYC1b}$ gpa1(468-472)(KIGII>QYELL) | Kapolka et al., 2020 | DI DCyFIR P1 S SSTR5 | somatostatin receptor type 5 | SSTR5 |
| BY4741 far1Δ sst2Δ ste2Δ fig1Δ::mTq2 X-2:$P_{TEF1a}$-SUCNR1-$T_{CYC1b}$ gpa1(468-472)(KIGII>ECGLY) | Kapolka et al., 2020 | DI DCyFIR P1 I SUCNR1 | succinate receptor 1 | SUCNR1 |
| BY4741 far1Δ sst2Δ ste2Δ fig1Δ::mTq2 X-2:$P_{TEF1a}$-SUCNR1-$T_{CYC1b}$ gpa1(468-472)(KIGII>GCGLY) | Kapolka et al., 2020 | DI DCyFIR P1 O SUCNR1 | succinate receptor 1 | SUCNR1 |
| BY4741 far1Δ sst2Δ ste2Δ fig1Δ::mTq2 X-2:$P_{TEF1a}$-SUCNR1-$T_{CYC1b}$ gpa1(468-472)(KIGII>DCGLF) | Kapolka et al., 2020 | DI DCyFIR P1 T SUCNR1 | succinate receptor 1 | SUCNR1 |
| BY4741 far1Δ sst2Δ ste2Δ fig1Δ::mTq2 X-2:$P_{TEF1a}$-SUCNR1-$T_{CYC1b}$ gpa1(468-472)(KIGII>YIGLC) | Kapolka et al., 2020 | DI DCyFIR P1 Z SUCNR1 | succinate receptor 1 | SUCNR1 |

| | | | | |
|---|---|---|---|---|
| BY4741 far1Δ sst2Δ ste2Δ fig1Δ::mTq2 X-2:$P_{TEF1a}$-SUCNR1-$T_{CYC1b}$ gpa1(468-472)(KIGII>EYNLV) | Kapolka et al., 2020 | DI DCyFIR P1 Q SUCNR1 | succinate receptor 1 | SUCNR1 |
| BY4741 far1Δ sst2Δ ste2Δ fig1Δ::mTq2 X-2:$P_{TEF1a}$-SUCNR1-$T_{CYC1b}$ gpa1(468-472)(KIGII>EFNLV) | Kapolka et al., 2020 | DI DCyFIR P1 14 SUCNR1 | succinate receptor 1 | SUCNR1 |
| BY4741 far1Δ sst2Δ ste2Δ fig1Δ::mTq2 X-2:$P_{TEF1a}$-SUCNR1-$T_{CYC1b}$ gpa1(468-472)(KIGII>EINLL) | Kapolka et al., 2020 | DI DCyFIR P1 15 SUCNR1 | succinate receptor 1 | SUCNR1 |
| BY4741 far1Δ sst2Δ ste2Δ fig1Δ::mTq2 X-2:$P_{TEF1a}$-SUCNR1-$T_{CYC1b}$ gpa1(468-472)(KIGII>DIMLQ) | Kapolka et al., 2020 | DI DCyFIR P1 12 SUCNR1 | succinate receptor 1 | SUCNR1 |
| BY4741 far1Δ sst2Δ ste2Δ fig1Δ::mTq2 X-2:$P_{TEF1a}$-SUCNR1-$T_{CYC1b}$ gpa1(468-472)(KIGII>QLMLQ) | Kapolka et al., 2020 | DI DCyFIR P1 13 SUCNR1 | succinate receptor 1 | SUCNR1 |
| BY4741 far1Δ sst2Δ ste2Δ fig1Δ::mTq2 X-2:$P_{TEF1a}$-SUCNR1-$T_{CYC1b}$ gpa1(468-472)(KIGII>QYELL) | Kapolka et al., 2020 | DI DCyFIR P1 S SUCNR1 | succinate receptor 1 | SUCNR1 |

**Dataset S2. Names and pKa values for all ligands used in this study (Related to Figs. 2, 3, and 4).**

| Ligand | 10X vehicle | pK_1 | pK_2 | pKa type | pKa source |
|---|---|---|---|---|---|
| platelet activating factor | 4% BSA | 1.86 | | predicted | DrugBank (ChemAxon) |
| kynurenic acid | water | 3.17 | | predicted | DrugBank (ChemAxon) |
| succinate | water | 4.21 | 5.64 | experimental | DrugBank (ChemAxon) |
| acetate | water | 4.54 | | predicted | DrugBank (ChemAxon) |
| niacin | water | 4.75 | | experimental | DrugBank (ChemAxon) |
| butyrate | water | 4.82 | | experimental | DrugBank (ChemAxon) |
| [Arg]8-vasopressin | water | 7.65 | 11.5 | predicted | DrugBank (ChemAxon) |
| epinephrine | 10 mM sodium ascorbate | 8.59 | | experimental | DrugBank (ChemAxon) |
| dopamine | 10 mM sodium ascorbate | 8.93 | | experimental | DrugBank (ChemAxon) |
| serotonin | water | 9.31 | 10 | predicted | DrugBank (ChemAxon) |
| isoprenaline | 10 mM sodium ascorbate | 9.81 | 8.96 | predicted | DrugBank (ChemAxon) |
| HU-210 | 20% MeOH | 9.98 | | predicted | DrugBank (ChemAxon) |
| adenosine | water | 12.45 | 3.94 | predicted | DrugBank (ChemAxon) |
| caffeine | water | 14 | | experimental | DrugBank (ChemAxon) |
| melatonin | 10% EtOH | 15.8 | | predicted | DrugBank (ChemAxon) |
| prostaglandin E2 | 4% BSA | 4.3 | | predicted | DrugBank (ChemAxon) |
| sphingosine-1-phosphate | 4% BSA | 1.51 | 9.7 | predicted | HMDB (ChemAxon) |
| 3-hydroxyoctanoic acid | 1% EtOH | 4.84 | | predicted | HMDB (ChemAxon) |
| lysophosphatidic acid | 4% BSA | 1.5 | | predicted | YMDB (ChemAxon) |

| | | |
|---|---|---|
| 2-arachidonoylglycerol | 4% BSA | not reported |
| JWH-018 | 20% MeOH | not reported |
| SRIF-14 | water | not reported |
| ZM-241385 | 10% EtOH | not reported |
| SCH-58261 | 10% DMSO | not reported |
